# Dive Performance and Aquatic Thermoregulation of the World’s Smallest Mammalian Diver, the American Water Shrew (*Sorex palustris*)

**DOI:** 10.1101/2021.06.02.446801

**Authors:** Roman W. Gusztak, Robert A. MacArthur, Kevin L. Campbell

**Affiliations:** Department of Biological Sciences, University of Manitoba, Winnipeg, Manitoba R3T 2N2 Canada; Department of Anesthesiology, University of Saskatchewan, Saskatoon, Saskatchewan S7N 0W8 Canada

**Keywords:** aerobic dive limit, aquatic thermoregulation, body oxygen store, diving behavior, energetics, hemoglobin, lung volume, muscle buffering capacity, myoglobin, Soricidae

## Abstract

Allometry predicts that the 12–17 g American water shrew (*Sorex palustris*)—the world’s smallest mammalian diver—will have the highest diving metabolic rate coupled with the lowest total body oxygen storage capacity, skeletal muscle buffering capacity, and glycolytic potential of any endothermic diver. Consistent with expectations, and potentially owing to their low thermal inertia, water shrews had a significantly higher diving metabolic rate in 10°C (8.77 mL O_2_ g^−1^ hr^−1^) compared to 30°C water (6.57 mL O_2_ g^−1^ hr^−1^). Unlike larger-bodied divers, muscle myoglobin contributed minimally (7.7–12.4%) to total onboard O_2_ stores of juvenile and adult water shrews, respectively, but was offset by high blood O_2_ carrying capacities (26.4–26.9 vol. %). Diving was predominantly aerobic, as only 1.2–2.3% of dives in 10 and 30°C water, respectively, exceeded the calculated aerobic dive limits at these temperatures (10.8–14.4 sec). The mean voluntary dive time of water shrews during 20-min trials in 3–30°C water was 5.0±0.1 sec (*N*=25, *n*=1628), with a mean maximum dive time of 10.1±0.4 sec. However, the average dive duration (6.9±0.2 sec, *n*=257) of radio-telemetered shrews exclusively foraging in a simulated riparian environment (3°C water) for 12- to 28-hr suggest that mean (but not maximum) dive times of water shrews in the wild may be longer. Mean dive duration, duration of the longest dive, and total time in water all decreased significantly as water temperature declined, suggesting that shrews employed behavioral thermoregulation to defend against immersion hypothermia. Additionally, free-diving shrews in the 24-hr trials consistently elevated core body temperature by ∼1°C immediately prior to initiating aquatic foraging bouts, and ended these bouts when body temperature was still at or above normal resting levels (∼37.8°C). We suggest this observed pre-dive hyperthermia aids to heighten the impressive somatosensory physiology, and hence foraging efficiency, of this diminutive predator while submerged.

## Introduction

The three primary factors dictating the maximum underwater endurance of air-breathing mammalian divers—anaerobic capacity, total onboard oxygen stores, and the rate at which these reserves are consumed—are all strongly influenced by body size (Emmett and Hochachka 1981; Schreer and Kovaks 1997; Halsey et al. 2006). The mass-specific rate of O_2_ consumption (V̇ O_2_), for instance, varies inversely with body mass according to a scaling coefficient of –0.25 to –0.33 (Kleiber 1975; White and Seymour 2003). Conversely, the catalytic activity of glycolytic enzymes (i.e., glycogen phosphorylase, pyruvate kinase, and lactate dehydrogenase), and hence the maximum anaerobic potential of muscle, scales positively with size among mammals with an exponent of 1.09 to 1.15 (Emmett and Hochachka 1981). Finally, body oxygen stores representing the combined oxygen reserves of lungs, blood, and muscle scale approximately isometrically with body mass (Calder 1996). Taken together, the resultant elevation in mass-specific V̇ O_2_ of small amphibious species actively swimming underwater (diving metabolic rate, DMR), coupled with lower absolute oxygen stores, should lead to a disproportionate reduction in their aerobic dive performance relative to larger divers. Maximal submergence times should be further curtailed by a reduced ability to exploit anaerobic pathways for ATP production and buffer aerobic (CO_2_) and anaerobic byproducts (lactic acid). As with dive endurance, the body surface area-to-volume ratio varies inversely with body size, resulting in lower mass-specific thermoregulatory costs for larger divers compared to their smaller counterparts (MacArthur 1989). Small-bodied divers also have a limited capacity for enhancing pelage or tissue insulation, while large divers are often endowed with a thick blubber layer or fur that dramatically increases whole-body insulation (Favilla and Costa 2020). All of these factors predispose small, semi-aquatic species to higher mass-specific rates of heat loss in the aquatic medium than is the case for their larger counterparts.

Given the significant physiological challenges facing small-bodied amphibious divers, it is surprising that relatively few studies have examined the diving capacity and aquatic thermal biology of mammals weighing less than 5 kg (e.g., Dawson and Fanning 1981; MacArthur 1984a; Evans et al. 1994; Hindle et al. 2006; Harrington et al. 2012; Jordaan et al. 2021). Moreover, to our knowledge, comparable data for the smallest mammalian divers (<100 g) are available only for star-nosed moles, *Condylura cristata* (McIntyre et al. 2002), European water shrews, *Neomys fodiens* (Köhler 1991; Vogel 1998), and, to a more limited extent, American water shrews, *Sorex palustris* (Calder 1969; McIntyre 2000).

Weighing in at 12 to 17 g, American water shrews are the world’s smallest endothermic divers. Well adapted to the cold, *S. palustris* primarily exploit riparian habitats within the boreal forests of North America, and can be found from Labrador in the east to above 60°N in the Northwest Territories, Yukon, and Alaska in the west (Whittaker et al. 2008). Semi-aquatic in nature, they forage both in and along the edges of ponds and fast-flowing streams and rivers, with up to 50–80% of their food intake coming from minnows, tadpoles, insect larvae, nymphs, crayfish, and other aquatic invertebrates (Conaway 1952; Sorenson 1962; but see Hamilton 1930). Somewhat surprisingly, vision plays a very limited role for underwater food detection (Svihla 1934) with prey capture success shown to be unaffected in total darkness (Catania et al. 2008). Instead, this flush-pursuit predator relies heavily on its remarkably acute nasal vibrissae to locate prey (even remotely through water movements), which they then attack with astonishing speed (<50 msec). The added ability to identify sedentary prey underwater via nasal olfaction further cements American water shrews as one of the most adept and agile underwater hunters on the planet (Catania et al. 2008; Catania 2013).

Consistent with their aggressive predatory lifestyle, *S. palustris* are able to consume >10% of their body mass in a single sitting and, accordingly, have the highest mass-specific BMR of any eutherian diver examined to date (∼3 times the mass-predicted value for a similar-sized mammal; Gusztak et al. 2005). Their strong positive buoyancy additionally requires constant paddling to remain submerged (Svihla 1934), and further suggests that the water shrew may also have an extremely high DMR and thus be limited to very short dives if it routinely stays within its calculated aerobic dive limit (cADL). Due to its small thermal inertia and large relative surface area, the water shrew should also lose heat rapidly while swimming and diving. As with other soricine (red-toothed) shrews, *S. palustris* are active year-round, though presumably are faced with greater energetic costs than terrestrial shrews given their aquatic lifestyle. This is especially true during winter while foraging under the ice, when convective heat loss is likely to pose a formidable challenge. Consequently, this diminutive insectivore is of interest not only in terms of its dive endurance, but also with respect to its thermoregulatory competence since, in theory, it should be highly susceptible to immersion hypothermia after even brief periods of aquatic activity.

The primary aim of this study was to investigate the dive performance and aquatic thermoregulatory ability of wild-caught *S. palustris* in both laboratory and semi-natural settings. Briefly, 20-min aquatic trials were completed to assess the diving behavior of this species, as well as test for the occurrence of behavioral thermoregulation in water shrews voluntarily diving over a range of water temperatures. To provide insight into the extent to which water shrews depend on aerobic metabolism while submerged, a second major objective was to determine this species’ cADL, which required measuring DMR and determining its total body oxygen storage capacity. Finally, to test whether this species takes advantage of immersion hypothermia to extend its dive duration while foraging in cold water, radio-implanted water shrews were provided access to a semi-natural riparian environment (water temperature=3°C) where they were required to forage exclusively under water during 12- to 28-hr trials.

## Methods

### Animal Capture and Care

Sixty-seven water shrews were live trapped in Whiteshell (49°47′N, 95°13′W) and Nopiming (50°28′N, 95°15′W) Provincial Parks, Manitoba, Canada, using Sherman live traps (256 cm × 76 cm × 76 cm). For comparative analysis, 18 sympatric though strictly terrestrial (20-30 g) short-tailed shrews, *Blarina brevicauda*, were also captured. Trapping techniques and holding conditions in captivity are detailed elsewhere (Gusztak and Campbell 2004; Hindle et al. 2009). Briefly, traps baited with frozen minnows were set out overnight and inspected every 2 hr to minimize trap mortality. Water shrew traps were placed on the edges of streams and ponds with abundant sedge, while short-tailed shrews were trapped in grass thickets located a few meters inland from the water shrew sets. Immediately upon capture, shrews were placed individually into covered 38-l plastic containers fitted with screen lids and supplied with soil, dried grass and leaves, a layer of thick moss, rocks, logs, a nest box (100 mm × 100 mm × 125 mm), and water trays. Provisions included meal worms, *Tenebrio molitor*, hulled sunflower seeds, ground Purina cat food^™^, and a prepared meat mixture (see below), along with any invertebrates found while trapping. Shrews were transported to the University of Manitoba Animal Holding Facility on the morning following capture.

Vogel (1990) suggested that captive European water shrews lose the hydrophobic properties of their pelage when animals are not provided with holding conditions that permit diving and access to dry moss for burrowing following aquatic activity. Consequently, much care was taken in the design and continued maintenance of the holding tanks to ensure the fur of water shrews was always in optimal condition. Modified 264-l glass aquaria (88 cm × 50 cm × 60 cm) with screened lids served as individual long-term holding containers for water shrews (see Gusztak and Campbell 2004). Each holding tank had a discrete terrestrial (∼75%) and aquatic (∼25%) compartment separated by a 1-cm thick Plexiglas partition. Short-tailed shrews were individually housed in 76-l terrestrial containers and were supplied with water dishes that were refilled every 12 hr. For both species, the terrestrial area was furnished as described above for the transport containers. Both species were offered a prepared meat mixture (beef and chicken hearts, pig and beef liver, ground beef, fish fillets, and canned dog food mixed with vitamin and calcium supplements) every 12 hr. To ensure water shrews were habitually swimming and diving in the setup, they were required to swim across the tank and dive under a removable partition to access the feeding tray. Additionally, mealworm larvae and aquatic prey (leeches, dragonfly nymphs, and small crayfish) were occasionally placed in the aquatic portion of the water shrew tanks to encourage natural foraging behavior. When offered, aquatic prey was preferentially consumed over the meat ration.

To minimize stress, shrews were never handled by hand. Transfer to and from the holding tanks was accomplished by placing a short (∼15 cm) blind-ending section of ABS tube (3.5 cm internal diameter) in the chamber, which the shrews readily entered. The tube was quickly capped and then opened following transfer, allowing the animal to exit freely. All wild shrews used in diving trials or for tissue processing were acclimated to holding conditions for a minimum period of three weeks before diving trials were initiated. Water shrews were allowed a recovery period of at least 48 hr between successive experimental trials (see below). As the integrity of the air boundary in the fur was found to strongly influence DMR, T_b_, and activity level, diving/metabolic trials in which the shrews fur became wetted were not included in any subsequent analyses.

Shrews are short-lived, with maximum life spans for both species in the wild being about 18 months (George et al. 1986; Beneski and Stinson 1987). Individual shrews were aged post-mortem based on the presence or absence of tooth growth rings following Hindle et al. (2009), and subsequently divided into juveniles (∼1–3 months of age) or adults (∼13–15 months). All study animals were captured under permission of Government of Manitoba Conservation trapping permits, and cared for in accordance with the principles and guidelines of the Canadian Council of Animal Care (University of Manitoba Animal Use Protocols: F01-025 and F05-014).

### Body Temperature Implants

Twelve water shrews were each equipped with model X-M transmitter (Mini-Mitter Inc., Sunriver, OR, USA) surgically implanted in the abdominal cavity. Transmitter mass (1.00–1.16 g) ranged from 6.7–9.7% of total body mass, and did not surpass the upper limit of 10% recommended by Brander and Cochran (1967). Five of these shrews underwent a second surgery 4 to 6 weeks later to implant replacement transmitters. Each transmitter was modified from the original packaging to decrease its overall size/mass and then calibrated following the method of Dyck and MacArthur (1992). Transmitter modification and surgical procedures are described in detail by Gusztak et al. (2005). Briefly, shrews were anesthetized with Isoflurane, first given at 3% for induction and then manually adjusted, as needed, between 2 and 3% to maintain a surgical plane of anesthesia. A midline incision through the skin and body wall was made along the *linea alba*. The sterilized transmitter was then placed into the abdominal cavity and incisions closed with sutures; no mortalities were recorded following this surgical procedure. Post-operative surgical care included placement of the shrews in a disinfected 38-l plastic container containing a nest box and shredded paper towel. Shrews were supplied with fresh food and water every 12 hr and were transferred back to their holding tanks after 48 hr. Aquatic trials started 7 to 10 days later.

### Voluntary Dive Behavior

We recorded the frequency and duration of voluntary dives by 25 water shrews (six of which were implanted with temperature-sensitive radio transmitters) over 20-min trials conducted in a 170.5 cm × 68 cm fiberglass dive tank filled with water to a depth of 60 cm (McIntyre et al. 2002). A transparent dive platform (17.5 cm × 68 cm) was situated at one end just above the waterline. A 4-cm section of ABS tube (internal diameter=3.5 cm) was fastened to the platform to provide a darkened refuge for the shrew. The center section of the tank contained open water (75 cm × 68 cm), while the remainder of the tank was covered with a sheet of 1-cm thick Plexiglas (78 cm × 68 cm) to encourage exploratory diving behavior. The Plexiglas sheet was equipped with handles so it could be removed quickly if a shrew became disorientated beneath.

Prey was absent from the dive tank. Instead, shrews were weighed and then placed in 38-l plastic containers supplied with soil, moss, logs and a nest box prior to the trials and offered a single mealworm 10 min before being transferred to the dive platform. To familiarize shrews with the dive arena, each animal completed a pre-trial training run in 30°C water. Behavioral dive trials were then conducted in randomized order in 3, 10, 20, or 30°C water. Trials were initiated when the shrew first entered the water and lasted precisely 20 min, upon which time the shrews were again transferred to the 38-l containers. Data were collected during the trial using a Sony^®^ microcassette recorder for post-trial analyses of dive durations and frequencies, time in water, and dive-surface ratios. For the latter, only post-dive surface intervals of <300 sec were included in the analysis.

### Body Temperature and Aquatic Thermoregulation

Rates of body cooling and re-warming were assessed for six implanted shrews during the above voluntary dive trials in 3–30°C water via body temperature (T_b_) data obtained during the 5-min pre-trial, 20-min dive trial, and 10-min post-trial periods, respectively. We also collected T_b_ profiles from eight implanted shrews placed in containers supplied with 1 L of 10, 20, and 30°C water, whereby they were allowed to swim or stand but not exit the water for the duration of the 10 min trials. To facilitate comparisons with the voluntary dive trials, rates of body cooling and re-warming were calculated for the first 5 min of the trial and post-trial periods, respectively. For both sets of experiments, radio signals from the Mini-mitters were continuously recorded via a Sony^®^ cassette recorder that was placed adjacent to the animal chamber/aquatic tank and T_b_ data subsequently analyzed at 1-min intervals.

### 24-Hour Dive Trials

In addition to the voluntary 20-min dive trials, relative activity, swim/dive times, and foraging behavior of one juvenile and four adult radio-implanted water shrews were continuously video recorded on a Sony^®^ Digital 8 video recorder in a simulated riparian environment maintained at 3°C for periods ranging from 12 to 28 hr. Body temperature was also recorded in parallel as described above and calculated at 1- and 4-min intervals during active and resting periods, respectively, and immediately before entering and after exiting the water. Each shrew was only tested once in this setup. For these trials, the previously described fiberglass tank was shortened to 128 cm and furnished with river-washed rocks and an artificial riverbank set embedded with tree roots and rocky crevices attached to one side (supplemental Fig. 1). The set had a ∼10 cm wide bank that transitioned from a dry surface to a depth of ∼2 cm and ran along the length of the aquatic enclosure. A 60-cm long clear PVC tube linked the water tank to a clear 64-l plastic terrestrial chamber so that the shrew was always visible when active. The terrestrial chamber was placed on a motion activity detector (MAD-1; Sable Systems Inc.) and furnished with a nest box, 2-3 cm of soil, and a 3-5 cm layer of fresh moss to promote tunneling behavior. Shrews were required to forage exclusively under water for invertebrate prey items (leeches, crayfish, mealworm larvae) which were replenished every 4 hr to ensure an abundant supply of food. Prior to data collection, water shrews were provided a 12-hr pretrial period to acclimate to the setup.

### Metabolic Cost of Diving

The costs of bouts of diving, grooming, and re-warming of 12 radio-implanted water shrews voluntarily diving in a 208 cm × 55 cm fiberglass tank filled with water to a depth of 44 cm were measured with open-flow respirometry following McIntyre et al. (2002). The tank was covered with three removable Plexiglas panels placed just below the waterline. A curved section of clear tubing 4 cm in diameter and 5 cm long connected the water tank to a 170-mL metabolic chamber constructed from a 6-cm length of Plexiglas tubing (internal diameter=6 cm). A removable rubber stopper at the rear of the chamber facilitated shrew transfer to and from the setup, while a removable partition was placed between the metabolic chamber and tank cover to prevent shrews from entering and exiting the water while pre- and post-dive metabolic measurements were recorded.

An outlet port was installed in the top/rear portion of the metabolic chamber while six inlet holes, each 1 mm in diameter, were drilled into the base of the chamber at the opposite end to facilitate mixing of air. Air was drawn sequentially at ∼500 mL min^−1^ from the chamber and through a column of Drierite using a TR-SS1 gas analysis sub-sampler (Sable Systems Inc., Las Vegas, USA) calibrated against a bubble flowmeter (accurate to within ±2%; Levy 1964). A subsample of this dry exhalent gas was analyzed using an Applied Electrochemistry S-3A O_2_ analyzer. Fractional O_2_ content was recorded at 1-sec intervals, while water (T_w_) and chamber temperature (T_a_) were recorded immediately prior to and following each trial. A respiratory quotient of 0.83 was assumed, based on an earlier study of fasted water shrews (Gusztak et al. 2005), and instantaneous V̇ O_2_ derived following the method of Bartholomew et al. (1981). Mean instantaneous V̇ O_2_ measurements were calculated at 20-sec intervals throughout the trial. Body temperature signals were recorded on a Sony^®^ cassette recorder and subsequently analyzed at 1-min intervals throughout the pre-trial, diving, and post-trial periods, respectively.

Concrete blocks and sections of ABS tubing were placed at the bottom of the tank to encourage longer exploratory dives, since preliminary trials suggested these objects increased time under water. Each shrew completed a total of four trials presented in random order: two in 10°C water and two in 30°C water. Trials conducted at 30°C provided an estimated DMR when thermoregulatory costs are presumably minimal (MacArthur 1989), while 10°C water was chosen to assess the thermoregulatory costs associated with submersion in cold water.

The mass of each shrew (corrected for telemeter mass) was recorded immediately before placing the animal in the metabolic chamber. Each trial consisted of a 10- to 15-min pre-trial period during which time the shrew was confined to the metabolic chamber and its lowest V· O_2_ over 5 min taken as the baseline value. After the partition was removed, 10 min was allotted for voluntary diving, which commenced upon the animal’s first entry into the water. At the end of the 10-min dive session, the partition was gently slid back into place to prevent further dives. The post-dive recovery V̇ O_2_ associated with re-warming and grooming was recorded until the animal’s V̇ O_2_ or T_b_ returned to within 95% of the pre-trial baseline. If this had not occurred after 15 min in the chamber, the trial was ended. Dive durations, grooming behavior, and relative activity were recorded on a Sony^®^ microcassette recorder and analyzed after each trial.

### Body Oxygen Stores

The total body oxygen stores of water shrews were determined after completing all diving/behavioral trials on each animal, while short-tailed shrews were allowed a 1- to 3-week acclimation period in the lab prior to determining their oxygen reserves. The mass of each shrew was recorded, after which the animal was deeply anesthetized with 3% Isoflurane inhalant anesthetic. A cardiac puncture was then performed to extract a blood sample for hemoglobin and hematocrit determinations (MacArthur 1984b; McIntyre et al. 2002), followed by euthanization via an overdose of Isoflurane. The heart, forelimb, and hindlimb muscles were then quickly removed and freeze-clamped in liquid nitrogen. Excised muscles were stored at –70°C and myoglobin content and buffering capacity determined later following the methods of Reynafarje (1963) and Castellini et al. (1981), respectively. The lungs were carefully removed after the majority of the heart muscle had been cut away, and lung volume determined gravimetrically following the procedures described by Weibel (1970/71). This involved inserting a 3-cm section of P20 cannula 5-8 mm into the trachea, with the cannula secured in place with a 5-0 silk ligature. Vetbond™ was applied to the knot at the juncture of the cannula/trachea to ensure the preparation would not slip. The trachea/lung prep was submerged in saline (0.9 M NaCl) and then inflated at a constant pressure of 20 mmHg with humidified air for ∼10–15 min before measurement. All volume measurements were corrected to STPD.

The total percentage of muscle mass, expressed as a fraction of digesta-free body mass, was also calculated for 12 water shrews and two short-tailed shrews. Skinned, eviscerated carcasses were submerged for ∼24–48 hr in a detergent solution at 32°C to detach any skeletal muscle adhering to the bones. The total skeletal muscle mass was then calculated by subtracting the bone mass from the initial carcass mass (MacArthur et al. 2001; McIntyre et al. 2002).

Total blood volume of water shrews was estimated from the allometric equation of Prothero (1980): blood volume (mL)=76 M^1.0^, where M=body mass in kg. Total body O_2_ stores (mL O_2_ in muscle, blood, and lungs, corrected to STPD) of *S. palustris* were determined following the procedures of Lenfant et al. (1970). This method assumes that water shrews dive with lungs fully inflated and with an initial lung oxygen concentration of 15%. The oxygen storage capacity of blood was calculated by assuming that 1/3 and 2/3 of the blood volume constituted the arterial and venous fractions, respectively, with the former having an oxygen saturation of 95% and the latter a 5% vol. decrease in O_2_ content compared to arterial blood (Lenfant 1970). Skeletal muscle myoglobin reserves were determined as the mean concentration of forelimb and hindlimb samples for each individual, multiplied by the estimated mass of skeletal muscle in the body. Blood and myoglobin oxygen capacities were presumed to equal 1.34 mL O_2_ g pigment^−1^ (Lenfant et al. 1970; Kooyman et al. 1983).

Total body oxygen stores were divided by the mean DMR (mL O_2_ sec^−1^) measured in 10 and 30°C water, in order to derive a cADL for water shrews diving at each T_w_. This estimate hinges on the assumption that all O_2_ stores are utilized during diving, before the animal switches to anaerobic respiration (Kooyman et al. 1980). We also calculated the behavioral ADL (bADL) at each T_w_, defined as the dive duration exceeded by only 5% of all voluntary dives (Kooyman et al. 1983).

### Skeletal Muscle Buffering Capacity

The skeletal muscle buffering capacities of short-tailed and water shrews were tested against non-bicarbonate buffers following the procedure of Castellini et al. (1981). Buffering capacity (slykes) was standardized to represent the μmol of base needed to increase the pH of 1 g of wet muscle mass from a pH of 6 to 7. A 0.3- to 0.5-g sample of frozen skeletal muscle comprised of a mixture of forearm and hindlimb tissue was ground up in 0.9 M NaCl, following which the solution was titrated with 0.2 M NaOH using an Accumet^®^ AB 15/15+ pH meter and an AccuTupH sensing electrode.

### Statistical Analyses of Data

For the 20-min voluntary dive trials we employed random slope, random intercept linear mixed models to assess the means of dive variables (i.e., mean dive duration, longest dive, five longest dives, total time in water, dive frequency, and log_10_ dive:surface ratio) across the four different water temperatures, with water temperature as the categorical variable and individual shrew as the random effect. These same models were also run for mean dive time and number of dives completed by radio-implanted shrews per five-minute interval, with latter modelled as the categorical variable and individual shrew as the random effect. Additionally, random intercept mixed models were run for each of the four water temperatures with pre-trial T_b_, lowest T_b_, mean trial T_b_, and highest post-trial T_b_ as the categorical variable and individual shrew as the random effect. Random intercept mixed models were also run for pre-trial T_b_, lowest T_b_, mean trial T_b_, highest post-trial T_b_, rate of T_b_ cooling, and total time in water with water temperature as the categorical variable and individual shrew as the random effect. Where applicable, all models were also run with water temperature as a continuous variable.

For the 10-min immersion trials, random intercept mixed models were run for T_b_ drop, rate of T_b_ cooling, and rate of T_b_ rewarming with water temperature as the categorical variable and individual shrew as the random effect. Finally, for the 24-hr trials, random slope, random intercept models were used to assess pre-dive T_b_ across the various time intervals, with both individual shrew and individual dives as random effects. This model was also run with time as a continuous variable, with the curvilinear regression being a better fit for the data than a linear fit.

All models were run using the function brm in the package brms (Bürkner 2017, 2018) in the R language and environment (R Core Team 2021). The package brms utilizes the program Stan (Carpenter et al. 2017). Bayesian p-values (hereafter denoted as ‘*p*’) were calculated using the function p_map in the package bayestestR (Makowski et al. 2019) in the R language and environment. Model residuals were visually checked for normality and heteroscedasticity, and log transformed as required.

Dive profiles were compared between implanted and non-implanted adult water shrews using the log likelihood ratio test (G-test, Zar 1974) and the means of variables compared using independent 2-tailed Student’s *t*-tests, as were the respiratory attributes of juvenile and adult water shrews, and adult short-tailed and adult water shrews. Regression lines for all other comparisons were fitted by the method of least squares. Significance for these analyses (hereafter denoted as ‘*P*’) was set at 5%. All means are presented as ±1 standard error (SE).

## Results

### Voluntary Dive Behavior

In most trials, water shrews were hesitant to dive until they had fully explored the surfaces of both the terrestrial and aquatic sections of the tank. Subsequently, predictable pre-dive behavior was routinely observed. Water shrews would approach and pause at the edge of the dive platform for 1–10 sec, during which time repetitive head nodding occurred, causing the shrew’s vibrissae to repeatedly touch the water. This behavior was typically followed by the shrew diving from the platform.

Water shrews engaged in two distinct categories of dives. Dives were classified as either shallow (<10 cm) or deep (reaching the tank bottom at 60 cm). Very few dives were completed between these depths, but if they occurred, were specified as shallow. No significant differences in diving duration or dive frequencies were found between juvenile and adult water shrews, nor between radio-implanted and non-implanted individuals (see below), and hence all data were pooled for subsequent analyses.

During each 20-minute trial, shrews completed, on average, 14.7±0.8 dives (range=0 to 50 dives). The mean dive time of 25 shrews (111 individual dive trials) was 5.0±0.1 sec (supplemental Fig. 2), with a median of 4.4 sec. Of the 1628 recorded voluntary dives, 322 (19.8%) were deep dives with an average duration of 8.0±0.2 sec. The five longest dives of each trial had a mean duration of 7.9±0.1 sec, while the mean longest dive per trial was 10.1±0.4 sec. Only three dives exceeded 20 sec with the longest voluntary dive recorded being 23.7 sec.

The average dive:surface ratio was 0.40±0.03. While short dives were often followed quickly by another dive, all dives >13 sec were accompanied by an extended (>30 sec) surface interval (supplemental Fig. 3). There was also a significant relationship between surface time and dive time, suggesting that longer dives require a longer recovery period than shorter dives (*p*<0.001, *r*^2^=0.042).

### Influence of Water Temperature on Dive Behavior

Eighteen shrews completed voluntary dive trials at all four water temperatures. For all dives combined, dive frequency was independent of T_w_ (*p*=0.33). However, T_w_ significantly influenced the total time water shrews spent swimming and diving (*p*<0.001), with shrews voluntarily spending just over half as long in 3°C water compared to 30°C water (Table 1). Mean dive duration increased with T_w_ (*p*<0.001; Fig. 1) while the dive:surface ratio decreased (*p*=0.049). On average, shrews surfaced for 68±5 and 53±4 sec before diving again in 3 and 30°C water, respectively. T_w_ also affected both the five longest (*p*<0.001) and single longest dive of each trial (*p*<0.001), with water shrews diving an average of 37% longer in 30°C compared to 3°C water (Fig. 1). The bADL of water shrews progressively increased from 8.8 sec in 3°C water to 12.2 sec in 30°C water.

**Figure 1.**
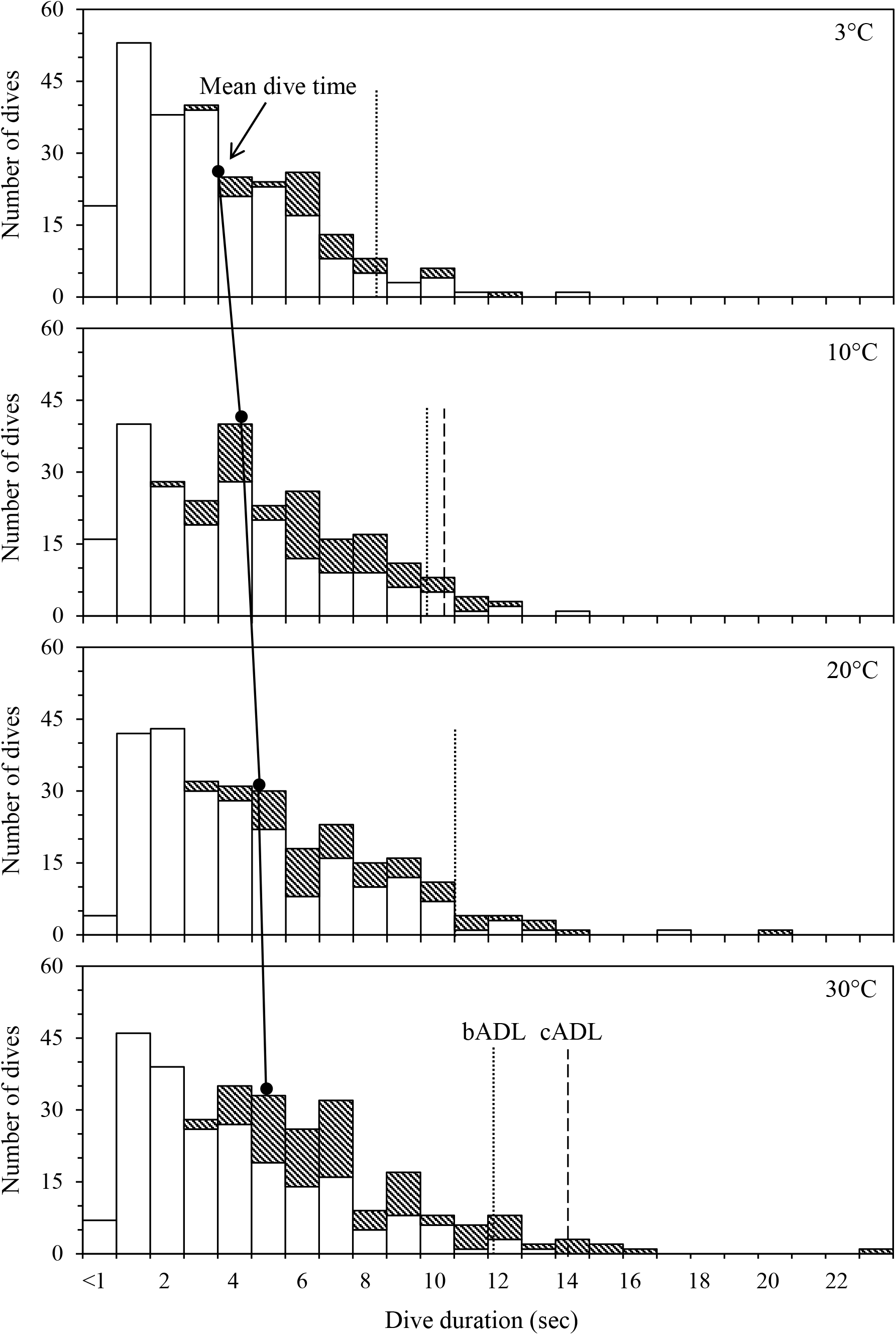
Influence of water temperature on the diving behavior of 18 American water shrews in 3–30°C water. Shallow dives (<10 cm deep) are denoted by open bars, while deep dives (60 cm) are denoted by stippled bars. Behavioral (bADL) and calculated aerobic dive limits (cADL; only calculated for 10 and 30°C) are denoted by dotted and dashed lines, respectively, while solid circles indicate the mean dive time at each water temperature.

**Table 1.** The effect of water temperature on the voluntary dive behavior (means ± 1 SE) of 18 captive American water shrews.

| Variable | Water temperature (°C) |  |  |  |
| --- | --- | --- | --- | --- |
|  | 3 | 10 | 20 | 30 |
| Total time in water (sec) | 114±10<sup>a</sup> | 125±14<sup>a</sup> | 169±15<sup>a,b</sup> | 217±21<sup>b</sup> |
| All dives (*n*) | 258 | 257 | 279 | 303 |
| Dive duration (sec) | 4.0±0.2<sup>a</sup> | 4.9±0.2<sup>a,b</sup> | 5.2±0.2<sup>b</sup> | 5.5±0.2<sup>b</sup> |
| Dive frequency (dives · min<sup>-1</sup>) | 0.72±0.08<sup>a</sup> | 0.71±0.09<sup>a</sup> | 0.78±0.07<sup>a</sup> | 0.84±0.11<sup>a</sup> |
| Dive:surface ratio | 0.29±0.09<sup>a</sup> | 0.38±0.05<sup>a,b</sup> | 0.48±0.07<sup>b</sup> | 0.42±0.04<sup>a,b</sup> |
| Duration of five longest dives (sec) | 6.3±0.3<sup>a</sup> | 7.6±0.3<sup>a,b</sup> | 8.5±0.3<sup>b</sup> | 9.3±0.3<sup>b</sup> |
| Duration of longest dive (sec) | 8.1±0.7<sup>a</sup> | 9.7±0.6<sup>a,b</sup> | 11.3±0.8<sup>b</sup> | 12.4±0.9<sup>b</sup> |

Note. Within each row, means sharing the same letter are not statistically different ( $P>0.05$ ).

### Influence of Transmitter Implants on Dive Performance

Dive performance was compared in twelve adult water shrews with (*N*=6) and without (*N*=6) an implanted 1.0-g abdominal temperature transmitter by first pooling the dive data for each group across all T_w_’s. There were no significant differences in dive performance between the two groups in any of the examined variables (supplemental Table 1). The frequency distribution of dives by implanted and non-implanted shrews was also compared for dives times binned by 3 sec intervals, using log likelihood ratios to ensure the calculated mean value was not influenced significantly by outliers (Supplemental Table 2). Again, no significant difference was found between implanted and non-implanted water shrews (*G* value=2.78, df=5, *P*>0.50).

### Body Temperature Profiles of Implanted Shrews

Twenty-three T_b_ data sets were obtained from six water shrews that completed 24 voluntary 20-min dive trials. Owing to a mechanical error with the recording device, no T_b_ data were obtained for one water shrew diving in 10°C water, though behavioral dive data were collected. The mean number of dives completed during the first 5 min was significantly greater than for any of the three subsequent 5-min periods (*p*<0.001), with ∼50% of voluntary dives occurring during the first 5 min of the trial (supplemental Fig. 4). T_b_ declined by ∼1.5°C during the initial 5-min period, with the rate of cooling not significantly affected by T_w_ (Table 2; Fig. 2). Following this initial curvilinear decline, T_b_ tended to plateau and was regulated near 38.8°C for the remainder of the dive trial. The lowest and mean recorded T_b’s_ during the diving session did not differ significantly between T_w_’s (*p*=0.49 and 0.55, respectively). Water shrews quickly elevated their T_b_ during the post-trial period, and attained a T_b_ similar to the pre-trial value regardless of T_w_ (Table 2; Fig. 2).

**Figure 2.**
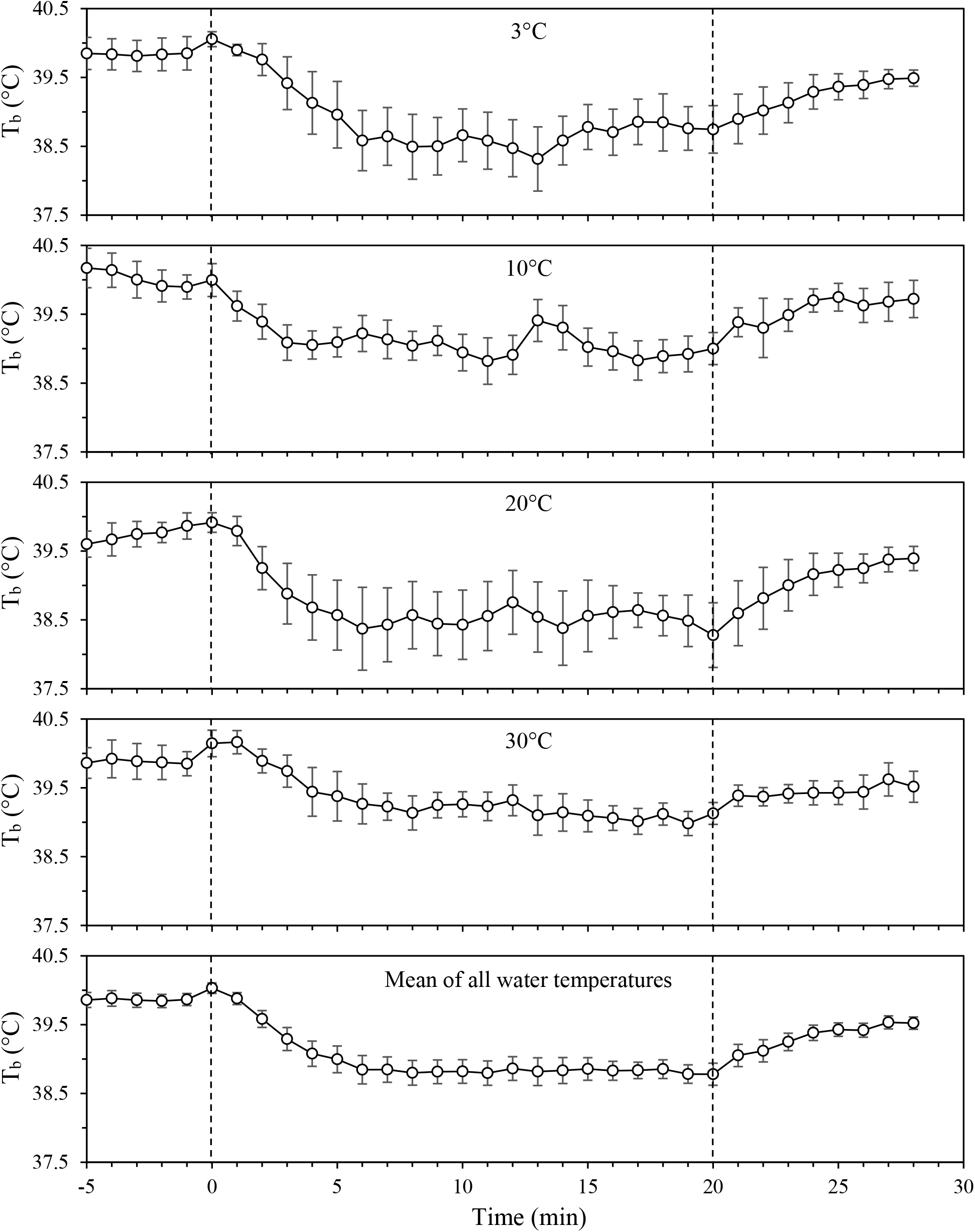
Telemetered body temperatures (T_b_) ± 1 SE of six American water shrews voluntarily diving during 20-min trials in 3–30°C water. Pre- and post-trial measurements were taken from shrews in temporary holding containers immediately before and after the trials (see text for details).

**Table 2.** Mean (± 1 SE) time in water, rate of cooling, and telemetered body temperatures (T_b_) recorded from six American water shrews during 20-min voluntary dive trials in 3–30°C water.

| Variable | Water temperature (°C) |  |  |  |
| --- | --- | --- | --- | --- |
|  | 3 | 10 | 20 | 30 |
| Time in water (sec) | 103±22<sup>a</sup> | 145±29<sup>a</sup> | 178±30<sup>a</sup> | 163±18<sup>a</sup> |
| Rate of cooling (°C min<sup>-1</sup>) | 0.22±0.09<sup>a</sup> | 0.18±0.04<sup>a</sup> | 0.27±0.07<sup>a</sup> | 0.15±0.08<sup>a</sup> |
| $T_b$ measurements (°C) | | | | |
| Pre-trial $T_b$ | 40.0±0.2<sup>a,1</sup> | 40.1±0.1<sup>a,1</sup> | 39.9±0.1<sup>a,1</sup> | 40.1±0.2<sup>a,1</sup> |
| Mean $T_b$ during diving | 38.9±0.3<sup>a,2,3</sup> | 39.1±0.2<sup>a,1,2</sup> | 38.6±0.3<sup>a,1,2</sup> | 39.4±0.2<sup>a,2</sup> |
| Lowest $T_b$ in trial | 38.0±0.3<sup>a,3</sup> | 38.4±0.2<sup>a,2</sup> | 37.7±0.4<sup>a,2</sup> | 38.8±0.2<sup>a,3</sup> |
| Highest post-trial $T_b$ | 39.5±0.2<sup>a,1,2</sup> | 39.8±0.2<sup>a,1</sup> | 39.8±0.2<sup>a,1</sup> | 39.5±0.1<sup>a,1,2</sup> |

Note. Rate of cooling measurements were determined from the first 5 min of each trial (see text for details). Superscript letters within a row sharing the same letter do not differ significantly ( $P>0.05$ ). Different superscript numbers within a column are significantly different ( $P>0.05$ ).

These results contrast with T_b_ profiles obtained from eight shrews that were prevented from exiting the water during 10-min trials in 10, 20, and 30°C water (supplemental Table 3; supplemental Fig. 5). Specifically, T_w_ significantly affected the overall T_b_ reduction, cooling rate, and rewarming rate, with shrews in 10°C water both cooling and rewarming 2.2 times faster than those in 30°C water.

### 24-Hour Dive Trials

Body temperature profiles and entry and exit times to and from the water were obtained from five shrews studied in the 12- to 28-hr trials. However, reliable submergence times were only obtained for three of the trials due to differences in camera position. While substantial variability was apparent, foraging activities were typically clustered into discrete bouts that consisted of 9.9±2.5 aquatic excursions and lasted 38.1±9.1 min each, with individual bouts separated by 65.7±8.8 min. Water shrews entered the water an average of 141 times per 24-hr period (range: 92-212) with a prey capture success rate of 28.6%, or ∼40 prey items per day. The shallow water of the riverbank shelf was consistently searched for prey before diving in the deeper water commenced, with shrews routinely capturing all prey items in the tank during each 4-hr period. Mealworms and crayfish were selectively consumed first, with excess food cached in a dedicated section of the terrestrial container.

The dive times of shrews actively foraging in the simulated riverbank enclosure were skewed towards longer duration dives (Fig. 3) than observed in the 20-min trials (Fig. 1). Consequently, the mean dive duration in the former set up (6.9±0.2 sec; *n*=227) was not only significantly longer than that of water shrews voluntary diving in 3°C water with no access to food (4.0±0.2 sec; *t*=11.42, df=483, *P*<0.0001), but also longer than the mean dive duration of dives completed during the 20-min dive trials (5.0±0.1 sec; *t*=8.33, df=1853, *P*<0.0001). Despite the right-shifted distribution, an abrupt drop in dive duration occurred between 9 and 11 sec yielding a bADL of 10.7 sec (Fig. 3).

**Figure 3.**
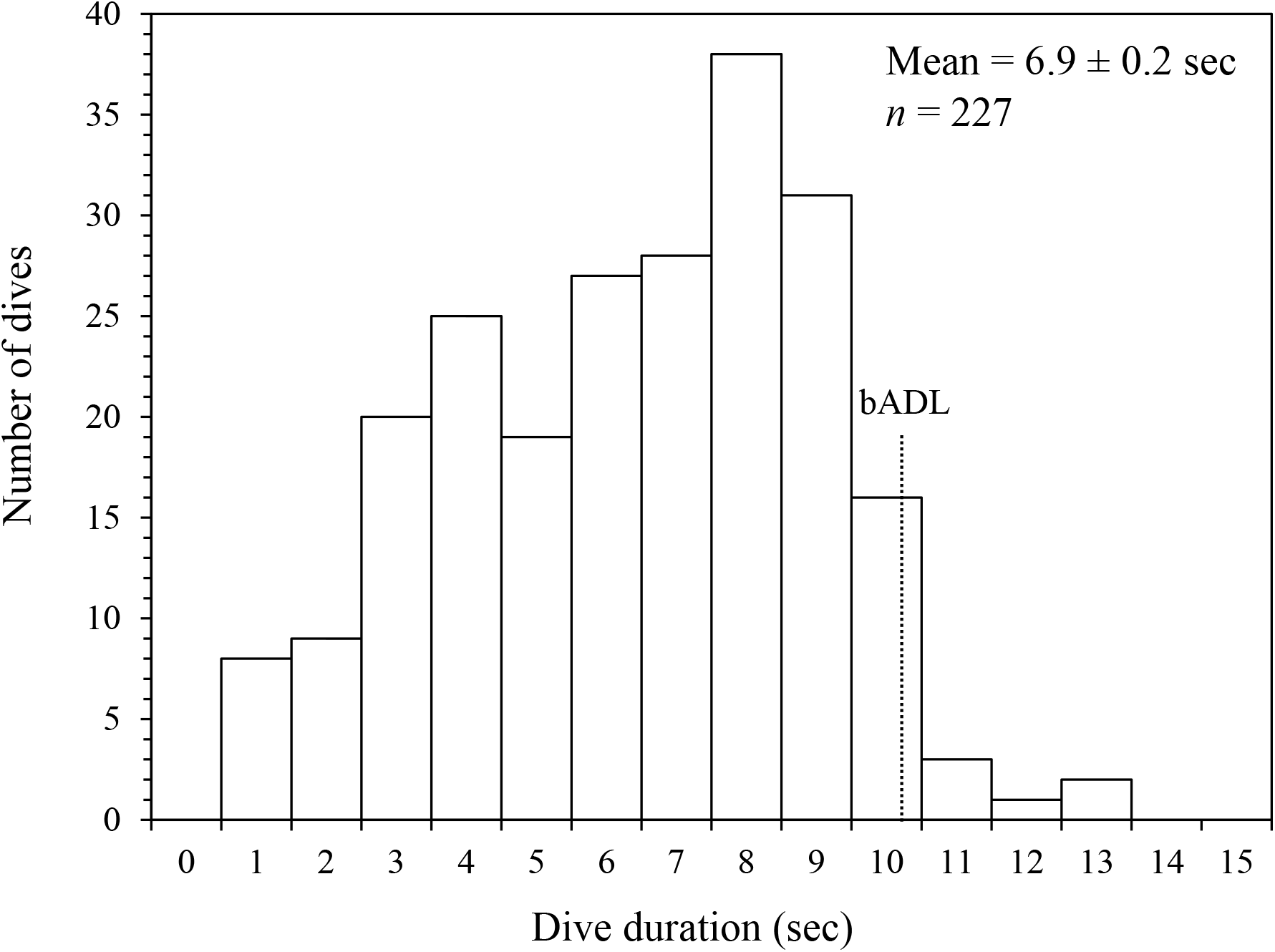
The distribution of dive times for three American shrews diving in a semi-natural riparian environment in 3°C water over 12- to 28-hr periods. The calculated behavioral aerobic dive limit (bADL) of 10.7 sec is indicated by the dotted line.

Body temperature profiles of free-ranging shrews exhibited several consistent patterns over the course of the trials. Principal among these was a significant (0.9±0.1°C) curvilinear elevation in T_b_ that initiated 8-12 min prior to initiating each aquatic foraging bout (Fig. 4). Consequently, mean T_b_ often averaged >38.8°C at the commencement of the first dive, though routinely dropped by 0.2–0.4°C upon exiting the water which was often followed by a further reduction in T_b_ (∼0.5 to 1.0°C). While shrews occasionally initiated another dive at T_b_’s <37.5°C, in most cases T_b_ was sharply increased again by re-warming in the terrestrial section prior to the next entry into water (Fig. 5 and supplemental Fig. 6). During the interbout intervals activity was minimal and core T_b_ was regulated near 37.5°C.

**Figure. 4.**
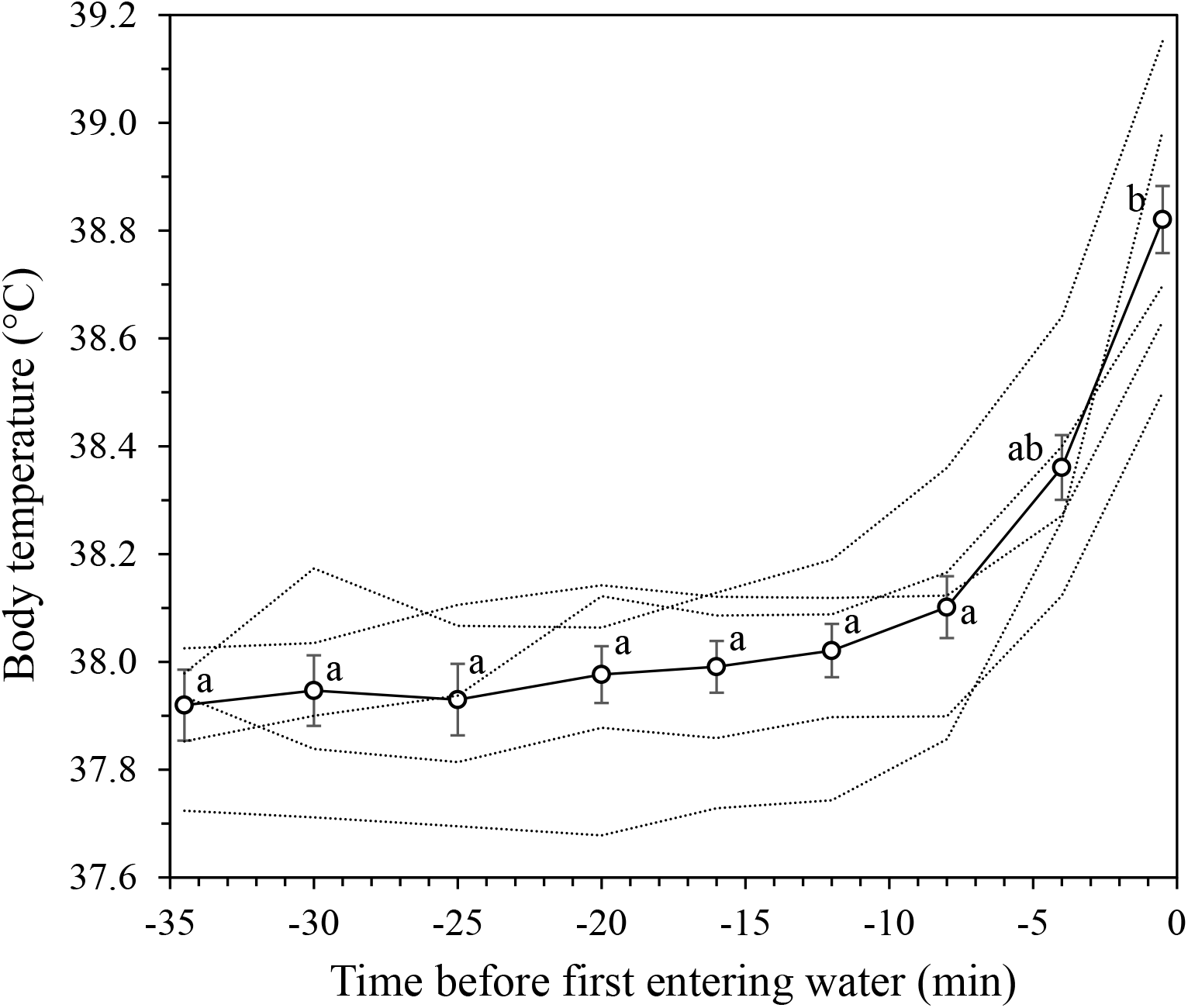
Overall mean (solid line; *n*=37) and individual mean (dotted lines) body temperature profiles (± 1 SE) of five American water shrews in the 35 min prior to initiating foraging bouts in a semi-natural riparian environment. Time points sharing the same letter are not statistically different (*P*>0.05).

**Figure 5.**
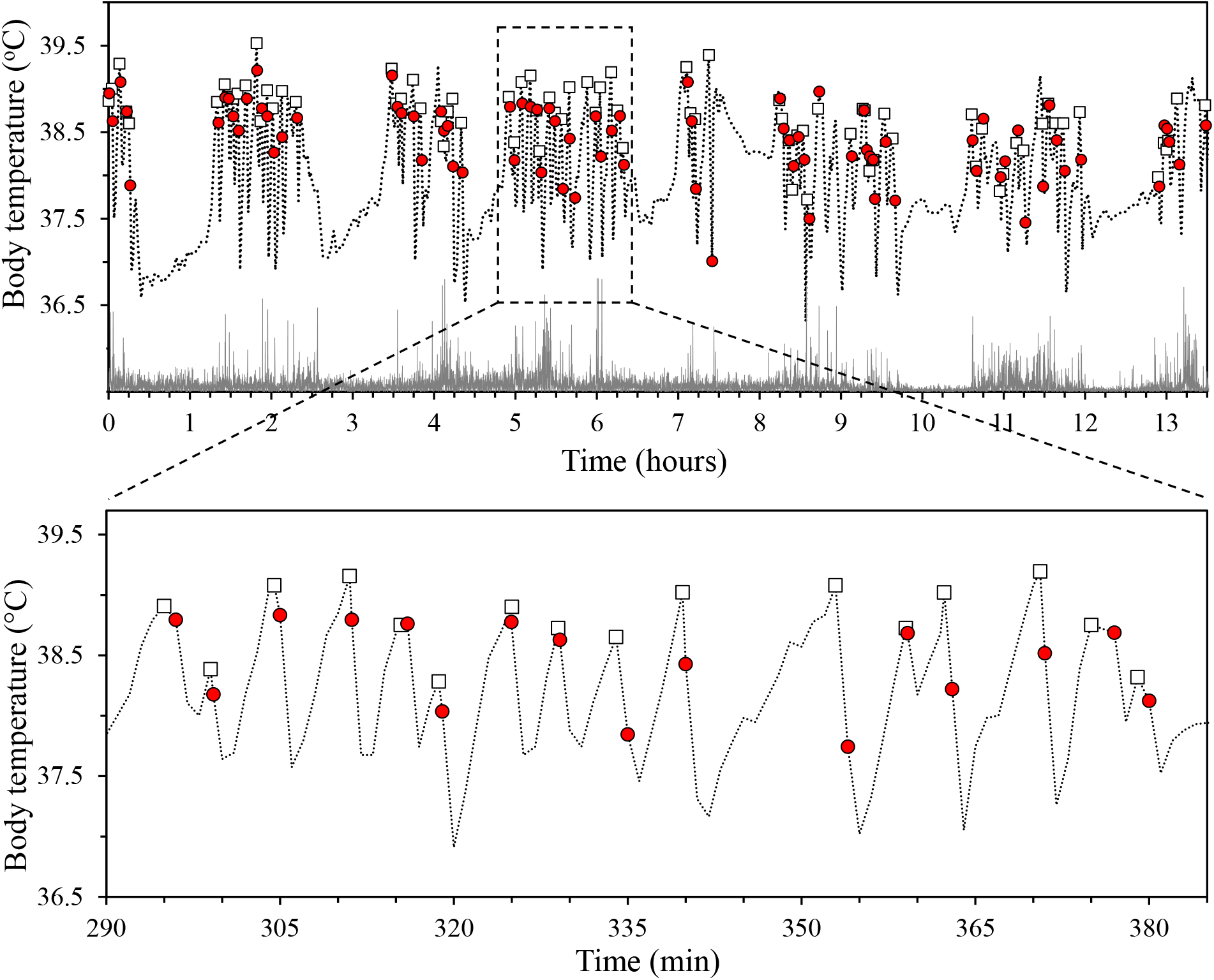
Body temperature fluctuations (dashed lines) and relative activity traces (grey bars) of an adult (13.3 g) American water shrew during a 13.5-hr trial in a natural riparian environment (water temperature=3°C). Open boxes represent body temperature at the start of each water excursion, while red circles denote body temperature immediately following exit from the water. Expanded inset box illustrates fine-scale body temperature profiles over the course of a single dive bout.

### Diving Metabolic Rate

The costs of repetitive diving and re-warming were determined for 12 implanted water shrews that completed 525 dives over 48 trials. Reliable T_b_ measurements were not obtained for 6 of the 48 trials, due to weak/absent signals from some transmitters. During the 10-min period available for voluntary diving, shrews spent an average of 51±7 sec (8.5% of total time) and 78±10 sec (13% of total time) diving in 10 and 30°C water, respectively (*t*=2.24, df=38, *P*=0.01). As with the behavioral dive trials (see above), mean dive time in 10°C water (3.8±0.2 sec; *n*=256) was significantly shorter than during the 30°C metabolic trials (5.4±0.3 sec; *n*=269) (*t*=2.54, df=32, *P*=0.007).

Integrity of the fur air boundary layer and level of activity in the metabolic chamber were both found to significantly affect T_b_ and increase recorded DMR estimates (data not shown). Consequently, we only utilized DMR values from water shrews whose pelage did not show evidence of wetting during the trial and that displayed minimal terrestrial activity. This resulted in estimates of DMR in only 13 of the 48 trials (five and nine from the 10°C and 30°C trials, respectively). Cumulative dive time per trial did not significantly influence DMR at either T_w_ (Fig. 6), though the mean DMR for shrews diving in 10°C water (8.77±0.30 mL O_2_ g^−1^ hr^−1^) was 1.33× greater than that in 30°C water (6.57±0.27 mL O_2_ g^−1^ hr^−1^).

**Figure 6.**
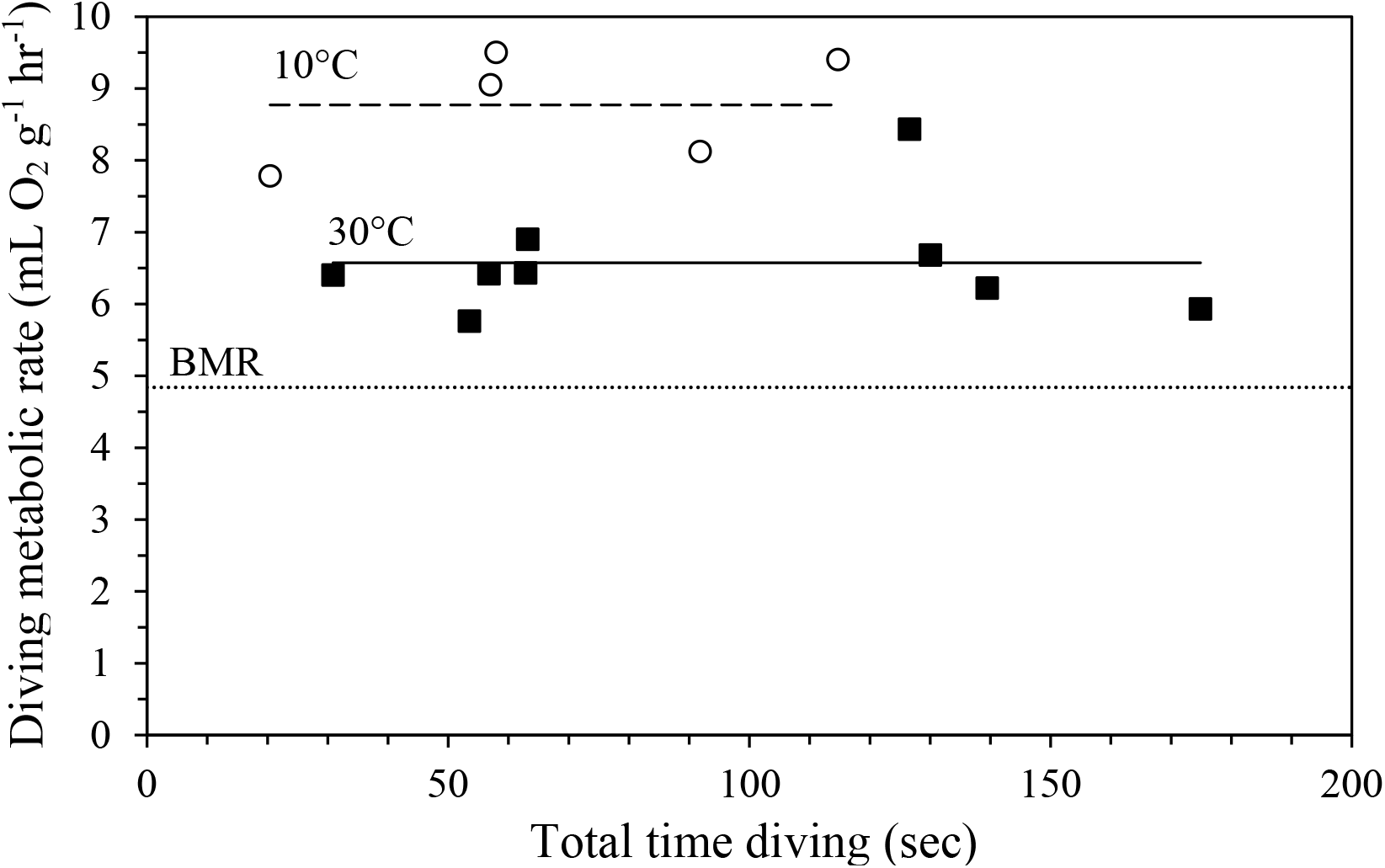
The relationship of diving metabolic rate to total submergence time of American water shrews voluntarily diving in 10°C (open circles, dashed line) and 30°C (closed squares, solid line). The dotted line denotes the basal metabolic rate (BMR; 4.84 mL O_2_ g^−1^ hr^−1^) of water shrews (Gusztak et al. 2005).

### Testing for Adaptive Hypothermia

Both regional heterothermy (peripheral hypothermy) and adaptive hypothermia have been forwarded as mechanisms that allow air breathing divers to extend their aerobic dive durations via temperature (Q_10_) induced reductions in metabolism (Favilla and Costa 2020). While it is unlikely that small-bodied water shrews can sustain meaningful peripheral heterothermy, reductions in core T_b_ may potentially allow them to extend their underwater endurance. Thus, voluntary dive times and concomitant core T_b_ were analyzed to test for potential linkages between these variables. We first plotted dive duration against T_b_ to determine if shrews with lower T_b_’s dove longer in cold water. To maximize the data set, dive data were combined for shrews completing voluntary dive trials in 3°C and 10°C water with those of shrews completing the DMR trials in 10°C water. A statistically significant negative relationship was found between dive duration and T_b_ (*r*^2^=0.0795, *P*<0.0001, *n*=309; Fig. 7a). Notably, shrews freely foraging in 3°C water in the semi-natural environment exhibited a similar trend (*r^2^*=0.0461, *P*=0.0103; Fig. 7b). However, a true measure of adaptive hypothermia should reflect reductions in core T_b_ being accompanied by a greater proportion of dive times above the cADL. Limiting these regression analyses to only voluntary dives that exceeded the predicted cADL at these temperatures (∼10 sec) did not return significant correlations (*r*^2^=0.002, *P*=0.801, *n*=27 and *r*^2^=0.001, *P*=0.871, *n*=28, respectively). Indeed, water shrews were reluctant to dive when cool, with few dives occurring at T_b_’s <37.5°C (Figs. 7c, d).

**Figure 7.**
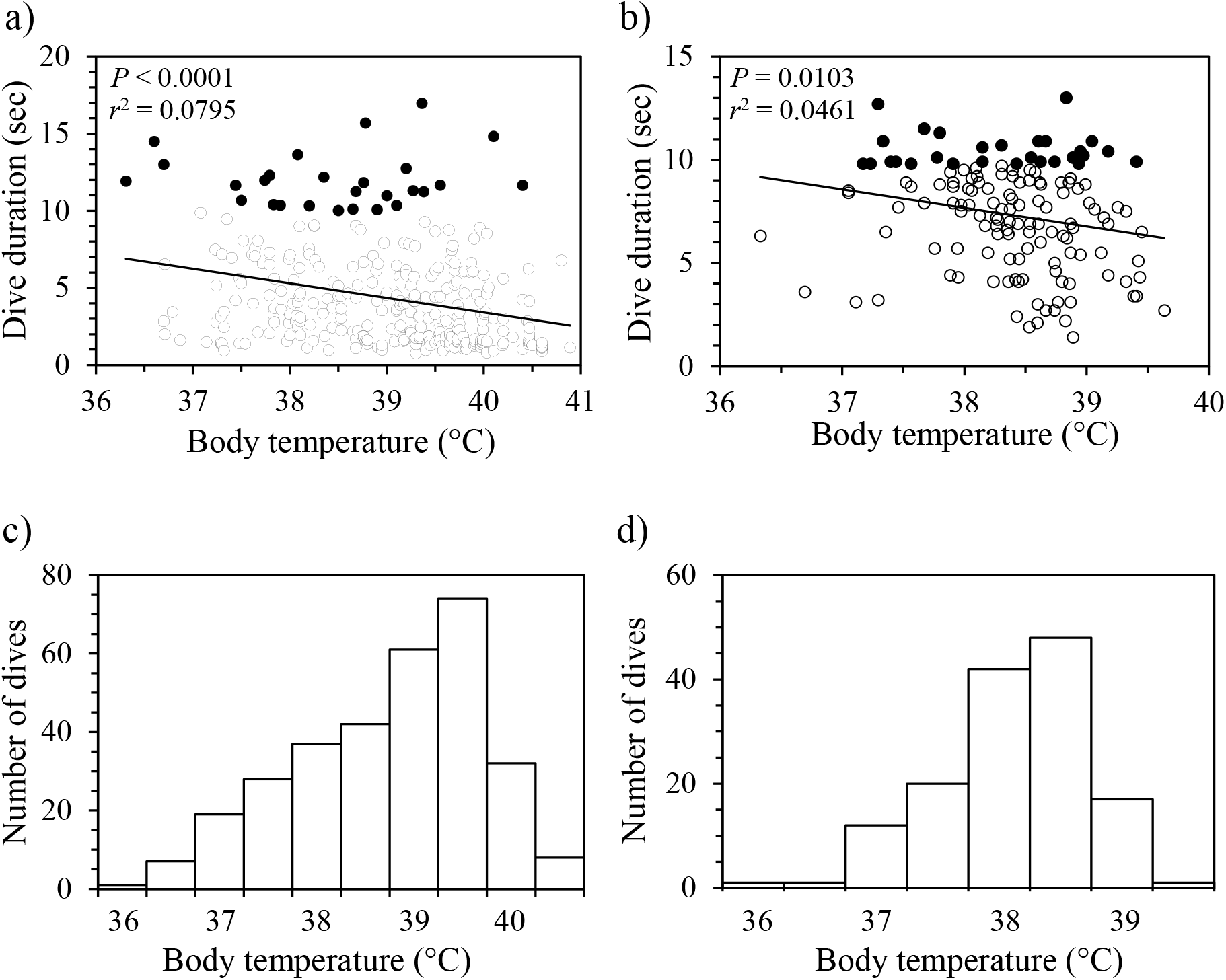
Dive duration and body temperature (T_b_) of American water shrews implanted with a ∼1.0-g intra-abdominal transmitter and voluntarily diving in a) 3°C and 10°C water (data pooled) and b) during the 12- to 28-hr trials in 3°C water. Although the relationship between dive duration and core T_b_ is significant in both instances, no significant relationship was observed when the analysis was limited dives greater than the calculated aerobic dive limit (∼10 sec; denoted by filled circles). The frequency distribution of T_b_ recordings in c) 3°C and 10°C water (data pooled) and d) during the 12- to 28-hr trials in 3°C water are also presented.

### Body Oxygen Stores and Muscle Buffering Capacity

Myoglobin concentration did not differ significantly between forelimb and hindlimb samples for either adults or juveniles in the two shrew species sampled (Table 3). The mean (forelimb and hindlimb) myoglobin concentration (mg g of wet tissue^−1^) of adult water shrews (6.04±0.25; *N*=9) was significantly higher than in juveniles (3.76±0.12; *N*=10; *t*=7.074, df=18, *P*<0.0001) and nearly 2× greater than for adult short-tailed shrews (3.03±0.12; *N*=9; *t*=7.624, df=17, *P*<0.0001). Skeletal muscle buffering capacity exhibited a similar trend, with adult water shrews having a higher value (38.22±2.28 slykes; *N*=13) than either juvenile water shrews (30.67±2.34 slykes; *N*=11; *t*=2.395, df=22, *P*=0.012) or adult short-tailed shrews (24.88±1.40 slykes; *N*=10; *t*=4.808, df=21, *P*<0.0001).

**Table 3.** Respiratory attributes of short-tailed and American water shrews.

| Variable | American water shrew |  | Short-tailed shrew |  |
| --- | --- | --- | --- | --- |
|  | Juvenile | Adult | Adult | Juvenile |
| Body mass (g) | 13.00±0.45 (20) | 14.57±0.36 (21)<sup>a</sup> | 24.26±1.41 (10)<sup>b</sup> | 21.78±0.13 (7) |
| Muscle mass as % of total body mass | 36.72±2.34 (2) | 40.51±3.38 (9) | 24.13±1.80 (2)<sup>b</sup> | No data |
| Forelimb Mb (mg g wet tissue<sup>-1</sup>) | 3.75±0.22 (10) | 6.18±0.31 (10)<sup>a</sup> | 2.98±0.32 (7)<sup>b</sup> | No data |
| Hindlimb Mb (mg g wet tissue<sup>-1</sup>) | 3.76±0.16 (10) | 5.89±0.45 (9)<sup>a</sup> | 3.01±0.40 (6)<sup>b</sup> | No data |
| Mean muscle Mb (mg g wet tissue<sup>-1</sup>) | 3.76±0.12 (10) | 6.04±0.25 (9)<sup>a</sup> | 3.03±0.26 (9)<sup>b</sup> | 2.70±0.49 (7) |
| Heart Mb (mg g wet tissue<sup>-1</sup>) | 10.97±1.22 (5) | 9.47±0.63 (6) | 8.77±0.55 (5) | 8.77±0.79 (5) |
| Buffering capacity (slykes)<sup>c</sup> | 30.67±2.34 (11) | 38.22±2.28 (13)<sup>a</sup> | 24.88±1.40 (10)<sup>b</sup> | 25.05±1.92 (6) |
| Total lung volume (mL STPD 100 g<sup>-1</sup>) | 4.55±0.18 (9) | 4.57±0.28 (5) | 3.34±0.07 (6)<sup>b</sup> | 3.43±0.11 (5) |
| Hematocrit (%) | 50.57±1.33 (10) | 50.23±1.33 (15) | 48.78±2.50 (8) | 50.17±2.50 (7) |
| Hemoglobin content (g 100 mL<sup>-1</sup>) | 19.69±1.02 (8) | 20.09±0.44 (13) | 17.29±0.62 (7)<sup>b</sup> | 16.87±1.54 (6) |
| Blood O<sub>2</sub> capacity (vol. %) | 26.38±1.37 (8) | 26.93±0.58 (13) | 23.18±0.84 (7)<sup>b</sup> | 22.61±0.14 (6) |

Note. Values are presented as means ± 1 SE, with number of animals sampled indicated in parentheses. Mb = myoglobin content.
<sup>a</sup>Values for adult and juvenile water shrews are significantly different ( $P < 0.05$ ).
<sup>b</sup>Values for adult water shrews and adult short-tailed shrews are significantly different ( $P < 0.05$ ).
<sup>c</sup>μmoles of base required to titrate the pH of 1 g of wet muscle by 1 pH unit.

Recorded lung volumes of adult (4.57±0.28 mL STPD 100 g^−1^; *N*=5) and juvenile (4.55±0.18 mL STPD 100 g^−1^; *N*=9) water shrews were similar (Table 3). These values are ∼1.1× greater than predicted by allometry (Stahl 1967) for a mammal of 15.2 g (adult) and 14.1 g (juvenile). Lung volumes of adult short-tailed shrews (3.34±0.07 mL STPD 100 g^−1^; *N*=6) were significantly less than those recorded for adult water shrews (*t*=7.401, df=9, *P*<0.0001) and 6% less than that predicted by allometry for a 22.8 g mammal.

Total blood oxygen capacity of adult and juvenile water shrews was high, averaging 26.93±0.58 and 26.38±1.37 vol. %, respectively (Table 3), with adult water shrews having a significantly higher blood oxygen capacity than adult short-tailed shrews (23.18±0.84 vol. %; *t*=0.896, df=9, *P*<0.0001). Hematocrit levels were high in both adult water shrews (50.23±1.33) and short-tailed shrews (48.78±2.50), and did not differ statistically between the two species (*t*=0.735, df=21, *P*<0.235). The calculated total O_2_ storage capacity of summer-caught adult and juvenile water shrews was 26.31 and 24.06 mL O_2_ STPD kg^−1^, respectively (Table 4). The mass-specific O_2_ storage capacity of summer-caught adult water shrews was 1.31× greater than that of adult short-tailed shrews (Table 4). The largest single contributor to total O_2_ stores in adults of both shrew species was O_2_ bound to hemoglobin in the blood, accounting for 62.5% and 68.5% in water shrews and short-tailed shrews, respectively.

**Table 4.** Oxygen storage capacities of the lungs, blood, and skeletal muscle of adult and juvenile American water shrews (*Sorex palustris*) and adult short-tailed shrews (*Blarina brevicauda*).

| Species | N | Oxygen stores (mL O<sub>2</sub> STPD kg<sup>-1</sup>) |  |  |  |  | Calculated ADL (sec) |  |
| --- | --- | --- | --- | --- | --- | --- | --- | --- |
|  |  | Lung | Arterial | Venous | Muscle | Total | Water temperature (°C) |  |
|  |  |  | blood | blood |  |  | 10 | 30 |
| Adult *S. palustris* | 8 | 6.60±0.08 | 6.33±0.04 | 10.12±0.07 | 3.26±0.21 | 26.31±0.24 | 10.8 | 14.4 |
| Juvenile *S. palustris* | 4 | 6.29±0.22 | 6.15±0.06 | 9.77±0.11 | 1.84±0.09 | 24.06±0.16 | 9.9 | 13.2 |
| Adult *B. brevicauda* | 2 | 5.01<sup>a</sup> | 5.45±0.37 | 8.37±0.75 | 1.32±0.04 | 20.14±1.07 | N/A | N/A |

Note. The aerobic dive limit (ADL) was calculated for a 14.0 g water shrew assuming a diving metabolic rate of 8.77 mL O<sub>2</sub> g<sup>-1</sup> hr<sup>-1</sup> in 10°C water and 6.57 mL O<sub>2</sub> g<sup>-1</sup> hr<sup>-1</sup> in 30°C water (see text for details).
<sup>a</sup>Value estimated from data in Table 3.

### Calculated Aerobic Dive Limit

Assuming that diving animals fully deplete their O_2_ reserves before initiating anaerobic respiration, and using the DMR estimate of 6.57 mL O_2_ g^−1^ hr^−1^ in 30°C water, adult and juvenile American water shrews have cADLs of 14.4 and 13.2 sec, respectively (Table 4). This value is consistent with the estimated bADL at 30°C (12.2 sec), determined from voluntary dive profiles (*n*=303 dives; Fig. 1). Due to an increased DMR in 10°C water (8.77 mL O_2_ g^−1^ hr^−1^), the cADL for adult and juvenile water shrews is 10.8 and 9.9 sec, respectively (Table 4). Again, these calculated limits closely match the bADL (10.2 sec) determined for water shrews diving in 10°C water (*n*=258 dives; Fig. 1). Only 1.2% and 2.3% of voluntary dives in 10°C and 30°C water, respectively, exceeded the cADLs for adult *S. palustris*.

## Discussion

As predicted by allometry, the diminutive size of *S. palustris* (mean=14.0 g) severely limits the species’ aerobic dive endurance. Indeed, adult water shrews exhibited the smallest total oxygen storage capacity (0.37 mL O_2_), the highest mass-specific DMR (6.57–8.77 mL O_2_ g^-1^ hr^-1^), and the lowest cADL (10.8–14.4 sec) of any mammalian diver studied to date. Moreover, water shrews have the highest surface-area-to-volume ratio of any endothermic diver, which should make them extremely susceptible to immersion hypothermia (MacArthur 1989). Given that aquatic foraging is one of the costliest means of feeding (Fish 2000), it is truly remarkable that this species is able to rely on this behavior to sustain its inherently high rate of heat production. This energetic burden is highlighted by the observation that captive American water shrews in a near thermoneutral (∼20°C) terrestrial setting must consume at least their body weight in prey on a daily basis (Gusztak et al. 2005). Hence, food requirements could conceivably be doubled or tripled in free-living individuals diving in ice-covered streams during the winter months.

### Body Oxygen Stores and Muscle Buffering Capacity

Divers are known for their considerable breath-hold capacity, a trait often attributed to disproportionately large body oxygen stores (Butler and Jones 1997; Favilla and Costa 2020). Not surprisingly, the mass-specific oxygen stores of water shrews were ∼30% greater than those of strictly terrestrial short-tailed shrews. Many researchers in this field consider the primary indicator of a diver’s breath-hold capacity to be its muscle oxymyoglobin concentration (Kooyman and Ponganis 1998, Ponganis et al. 1999; Mirceta et al. 2013). Adult *S. palustris* appears not to follow this trend as myoglobin accounts for only 12.4% of total body O_2_ stores, which is similar to alcids (∼5-10%), but substantially lower than values (∼30 to 50%) reported for various pinnipeds and toothed whales (Elliott et al. 2010; Favilla and Costa 2020). Moreover, skeletal muscle myoglobin concentration of adult water shrews (0.60 g 100 g^−1^; Table 3) is only half that recorded for other semi-aquatic mammals: star-nosed moles (1.36 g 100 g^−1^; McIntyre et al. 2002), muskrats (1.21–1.38 g 100 g^−1^; MacArthur et al. 2001), and beaver (1.2 g 100 g^−1^; McKean and Carlton 1977). Similar to other divers, however, myoglobin stores are relatively slow to develop as values for juveniles (<4-month old) were ∼62% those of adults and only accounted for 7.7% of total body O_2_ storage capacity.

The relatively low myoglobin concentrations in small-bodied alcids and semi-aquatic mammals likely arises from functional constraints imposed by muscle fiber types and relatively high mitochondrial volumes (Weibel 1985; Ordway and Garry 2004) that limit maximal attainable levels in muscle. Nonetheless, skeletal muscle myoglobin concentrations of adult water shrews are 2–4× those of strictly terrestrial short-tailed (0.16 to 0.30 g 100 g^−1^; Stewart et al. 2005 and Table 3) and Etruscan shrews, *Suncus etruscus* (0.15 g 100 g^−1^; Jürgens 2002), suggesting an adaptive increase associated with diving. This difference appears to be linked to the marked increase in the net surface charge of *S. palustris* myoglobin versus those of non-diving shrews, which is proposed to foster higher myoglobin concentrations by minimizing the potential for newly synthesized (apomyoglobin) and mature protein chains from aggregating/precipitating (Mirceta et al. 2013; Samuel et al. 2015; He et al. 2021). Of note, the highest myoglobin concentrations were found in the heart for both short-tailed (0.88 g 100 g^−1^) and water shrews (1.10 g 100 g^−1^). This finding presumably highlights a key role for this respiratory pigment in supporting the exceptionally high heart rates (>750 beats per minute) of soricids (Doremus 1965; Vornanen 1992), though this feature may also help extend aerobic metabolism of this tissue during long dives by water shrews.

The water shrew is able to compensate for its low skeletal muscle myoglobin concentration owing to the potential gain in O_2_ stores in the lungs and, especially, the blood. The mean lung volume of adult water shrews (4.57 mL STPD 100 g^−1^) was ∼1.1× greater than predicted by allometry for a 15.2 g mammal (Stahl 1967). Further, this species has a mass-specific pulmonary O_2_ storage capacity that is 1.14-1.37× greater than for adult short-tailed shrews (this study) and other shrew species studied to date (Gehr et al. 1981). The gain in lung volume of *S. palustris* could serve to increase buoyancy, as well as provide an important source of O_2_ while diving. It should also be noted that underwater sniffing plays an important role in food acquisition for this species (Catania 2006; Catania et al. 2008). In this context, the enlarged lungs of American water shrews may at least partially compensate for exhaled gas bubbles lost incidental to underwater sniffing (Gusztak, unpublished observations). However, lung volume of *S. palustris* is almost half that of the adult star-nosed moles (8.1 mL STPD 100 g^−1^; McIntyre et al. 2002), which also exploits underwater olfaction (Catania 2006). Another relevant observation is that submerged water shrews would routinely inspire trapped gas released from their fur that coalesced under the Plexiglas cover during the voluntary dive trials. This finding implies an ability to re-breath air bubbles trapped under the ice during winter foraging in order to extend submergence times as has been observed in muskrats (MacArthur 1992). Since water shrews commonly co-habit waterways with beavers and muskrats (Conaway 1952), it is expected that bubbles can similarly be exploited by *S. palustris* diving under the ice.

By far, the most important source of O_2_ for a diving water shrew is the large blood reserve, which comprises 62.5–66.2% of this diver’s O_2_ storage capacity. This is reflected in their high hemoglobin (19.9 g 100 mL^−1^) and hematocrit (50.4%) values, with the former resulting in a mean blood O_2_ capacity of 26.9 vol. % for an adult water shrew. Many species of shrews examined to date have high recorded hemoglobin (range: 15–19 g 100 mL^−1^) and hematocrit values (range: 45–57%), some of which are near the upper limits recorded for any terrestrial mammal (Sealander 1965; Wolk 1974; Gehr et al. 1981). Even so, the blood O_2_ capacity calculated for the American water shrew is the highest recorded for any soricid, including the European water shrew (23.9 vol. %; Wolk 1974), the Etruscan shrew (23.3 vol. %; Bartels et al. 1979) and adult short-tailed shrew (23.2 vol. %; Table 3). Adult water shrews also exhibited a higher blood O_2_ capacity than other small semi-aquatic divers studied to date, including adult star-nosed moles (23.0 vol. %; McIntyre et al. 2002) and muskrats (24.1 vol. %; MacArthur et al. 2001).

Buffering capacity of skeletal muscle has been shown to increase with increasing body mass and is also important for prolonging burst activity in species utilizing sprinting (Castellini et al. 1981; Hochachka and Mommsen 1983). In anaerobic or severely hypoxic conditions, as may be encountered during prolonged diving, glycolysis is the only, albeit inefficient, means of ATP production. This process is inhibited if intracellular pH drops too low, hence intracellular buffers are critical to ensure an optimum pH for glycolysis to occur while exercising in low O_2_ environments. Adult water shrews have a skeletal muscle buffering capacity (38.2 slykes) that is significantly greater (1.5-fold) than for adult short-tailed shrews. Although allometric scaling of glycolytic enzymes predicts that water shrews should have the poorest ability of any diver to utilize anaerobic glycogenolysis (Emmett and Hochachka 1981), adult water shrew buffer capacity was similar to platypus (38.2 slykes; Evans et al. 1994), star-nosed moles (44.1 slykes; McIntyre et al. 2002) and summer-caught adult muskrats (51.5 slykes; MacArthur et al. 2001). Since few voluntary dives by water shrews exceeded the cADL, glycolytic pathways probably do not play a large role in the majority of dives completed by this species. Instead, their elevated muscle buffering capacity presumably attenuates tissue pH changes arising from the rapid rate of CO_2_ accumulation during aerobic breath hold dives, though it may also be important for their flush-pursuit aquatic foraging strategy that involves periods of brief, intense motor activity (Catania et al. 2008).

### Diving Metabolic Rate

The American water shrew has the highest DMR recorded for any diving vertebrate. This trait reflects in part its inherently high mass-specific BMR (4.84 mL O_2_ g^−1^ hr^−1^; Gusztak et al. 2005), as well as the rapid limb strokes, high buoyancy, and possibly strong drag forces due to the species’ high mass-specific surface area. However, the DMR of *S. palustris* exposed to minimal thermal stress is only 1.4× BMR, which is less than star-nosed moles (2.1× RMR; McIntyre et al. 2002) and muskrats (2.7× BMR; MacArthur and Krause 1989), but similar to sea otters diving and capturing prey (1.6× RMR; Yeates et al. 2007).

### Dive Performance

Prior to this study, very little was known about the voluntary breath-hold capacity of *S. palustris*. Our captive water shrews exhibited a mean voluntary dive duration of 5.0 sec during the 20-min dive trials, which is close to the value (5.2 sec) reported by McIntyre (2000) for a single captive specimen, and is consistent with the ranges previously reported for *S. palustris* (1–15 sec; Svihla 1934; Calder 1968) and *N. fodiens* diving in captivity (3–6 sec; Churchfield 1985; Ruthardt and Schröopfer 1985). Vogel (1998) suggested that the dive times of captive water shrews were probably shorter than those of shrews in the wild due to tank size constraints and absence of aquatic prey. However, we believe that our study provides a reasonably accurate representation of the American water shrew’s dive capacity. For example, the maximum recorded dive time of 23.7 sec is 12% greater than that predicted for a 16.3 g diver (21.2 sec; Schreer and Kovacs 1997), but is virtually identical to the maximal dive time (24 sec) for a slightly larger European water shrew diving in a 2-m deep stream (Scholetch 1980). The mean dive time of this species is also slightly longer than predicted from allometry (cf., Fig. 4 of Jordaan et al. 2021). That said, dive times in the 20-min dive trials were likely skewed towards shorter mean (but not maximum) durations than expected in nature. Indeed, the mean submergence time of water shrews foraging in 3°C water in the artificial riverbank environment (6.9 sec) was ∼75% longer than that (4.0 sec) of shrews diving at the same temperature without access to food.

Even under semi-natural laboratory conditions, *S. palustris* is an impressive diver compared to other small-bodied divers. Another semi-aquatic insectivore, the star-nosed mole, is ∼4× larger (50–60 g), yet exhibits a mean dive duration that is only 1.3–1.8 times greater (9.2 sec; McIntyre et al. 2002). In contrast, most dives in the wild by the smallest (∼50 g) avian diver, the dipper (*Cinclus mexicanus*; <6 sec; Calder 1968), are shorter than for foraging water shrews, while the much larger 325–450 g bufflehead (*Bucephala albeola*) has a mean dive time of only 12.5 sec (Gauthier 1993). Even larger mink (*Mustela vison*; 850 g) and spotted necked otters (*Hydrictis maculicollis*; 4 kg) have comparable mean dive durations of 9.9 and 8.5 sec, respectively (Dunstone and O’Connor 1979; Jordaan et al. 2021). Finally, only moderately longer average dive times are observed for juvenile (300–500 g) and adult muskrats (650–900 g) (19.2–22.0 sec; MacArthur et al. 2001), and 1.5–2.0 kg free-ranging platypus, *Ornithorhynchus anatinus* (31.3 sec; Bethge et al. 2003).

Radio-equipped water shrews did not show any significant changes in dive performance compared to non-implanted animals, despite the transmitters being nearly 10% of their body mass. The strong buoyancy afforded by the large volume of air trapped in their pelage probably contributes to this lack of effect. In fact, the mass-specific air capacity (0.35 mL g^−1^) of the water shrew’s pelage is nearly double that of star-nosed moles (0.19 mL g^−1^; McIntyre 2000). American water shrews also have a lower specific gravity (0.761), or stronger buoyant force in water, compared to star-nosed moles (0.826; McIntyre 2000) and muskrats (0.952; MacArthur 1992), but less than that of European water shrews (0.726; Köhler 1991). While the marked buoyancy of *S. palustris* should increase the energetic costs of submerging to depth, their ability to rapidly and passively ascend the water column (Shilva 1934) should enable the retrieval of relatively large prey items while lessening the probability of their escape. Indeed, captive *S. palustris* were routinely observed passively surfacing with 2–3 g crayfish captured at the bottom of a 30-cm tank (Gusztak, unpublished observations). Likewise, Köhler (1991) recorded that *N. fodiens* could retrieve a 12.8 g lead-filled snail shell from the tank bottom.

### Aerobic Dive Limits and Diving Behavior

The relevance of determining cADL for certain diving species has been called into question since some large-bodied, benthic divers tend to routinely exceed their cADL. For instance, deep diving Australian and New Zealand sea lions often exceed their cADLs by 1.4 and 1.5-fold, respectively (Costa et al. 2001), while beaked whales may surpass their bADL by an astonishing 2.4 hrs (Quick et al. 2020). By contrast, small amphibious divers <2 kg submerge to relatively shallow depths with less interspecific variability in dive depth than larger species, and seldom dive beyond their cADL. For instance, the star-nosed mole (50 g) and muskrat (680 g) exceeded their cADLs during only 2.9% and 6% of all voluntary dives, respectively (MacArthur et al. 2001; McIntyre et al. 2002). In line with these observations, only 1.2–2.3% of dives by water shrews in 10–30°C water exceeded their cADLs. Similarly, dive duration frequencies exhibited an abrupt drop between 9 and 11 sec during the 24-hr trials, yielding a bADL (10.7 sec) that is virtually identical to the cADL of adult water shrews diving in 10°C water (10.8 sec). Taken together, these observations suggest a strict adherence to an aerobic diving schedule in this species. Thus, the cADL of small-bodied divers seems to provide a useful estimate of their breath-hold capacity, even though this is not always the case for larger divers.

### Influence of Water Temperature on Dive Behavior and Body Temperature

Presumably because it is energetically more efficient to prevent large drops in T_b_ than incur the post-immersion costs of re-warming, many small amphibious divers limit core T_b_ cooling behaviorally by periodically exiting the water to re-warm (MacArthur 1989; Hindle et al. 2006). Based on voluntary dive trials in 3–30°C water, shrews did not consistently show a statistically significant difference in any measured variable across adjacent T_w_’s. Instead, for most variables examined, there was a gradual positive relationship between the behavioral index of dive performance and increasing T_w_. Consistent with behavioral thermoregulation, large differences in dive performance between the two extreme T_w_’s were apparent. For example, relative to their performance in 3°C water, the same individuals in 30°C water extended mean and maximum dive durations by 1.3- to 1.5-fold while also increasing their deep dive frequency (3.3×) and total time spent in water (1.9×). While many other small-bodied semi-aquatic mammals and birds display a similar positive correlation of dive duration and frequency with T_w_ (MacArthur 1984a; de Leeuw et al. 1999; McIntyre et al. 2002), free-diving mink and European beaver do not (Harrington et al. 2012; Graf et al. 2018).

Regardless of water temperature, water shrews experienced the greatest drop in T_b_ during the first 5 minutes of each 20-min diving trial, which then plateaued near 38.5–39.0°C for the remaining 15 min. This latter interval was accompanied by a sharp reduction in aquatic activity, suggesting that water shrews principally rely on behavioral thermoregulation to limit further declines in T_b_ during bouts of aquatic activity. Similarly, free-ranging muskrats and beavers were observed to maintain core T_b_ within ±1°C of normothermic T_b_ throughout most semi-aquatic activity, and also appeared to use behavioral thermoregulation to limit body cooling (MacArthur 1979; Dyck and MacArthur 1992). In contrast to the voluntary dive trials when shrews were free to enter/exit the water, T_w_ significantly affected the rate of cooling during the 10-min immersion trials, with T_b_ declining nearly twice as much in 10°C vs. 30°C water (5.7°C vs. 2.9°C, respectively). Rates of cooling in 10°C water (0.91±0.06°C min^−1^) were, however, substantially lower than those previously measured by Calder (1969) for *S. palustris* at the same temperature: 2.06°C min^−1^ (swimming) and 2.46°C min^−1^ (diving). This difference may, in part, relate to our efforts to maintain the pelage of the shrews in optimal condition thereby preserving the integrity of the critical air layer trapped in the fur.

The adaptive hypothermia hypothesis (Butler and Jones 1997) has been advanced to explain why some divers routinely exceed their cADL. However, the benefit of increased aerobic dive endurance comes with the mandatory energetic cost of re-warming cooled tissues after diving and, for small-bodied divers especially, the potential impairment of locomotor function and foraging efficiency accompanying immersion hypothermia. Studies examining the adaptive hypothermia tenet have yielded mixed results (Ponganis et al. 2003; Hindle et al. 2006; Niizuma et al. 2007). Owing to their low O_2_ stores, high mass-specific DMR, and rapid rate of heat loss in water, water shrews might be expected to implement body cooling to prolong their underwater endurance. Consistent with this hypothesis, water shrews in dive trials at 3 and 10°C, and those freely foraging in 3°C water in the semi-natural environment, showed a statistically significant inverse relationship between dive duration and core T_b_. However, adaptive hypothermia should manifest as a greater proportion of dive times above the cADL in animals exhibiting the lowest core temperatures. No significant correlation was found between T_b_ and dive times that exceeded the cADL in either the acute or 24-hr dive trials. More telling was the finding that dive frequency significantly declined in concert with T_b_, with few dives occurring at T_b’s_ <37.5°C. This observation exemplifies the strict thermoregulatory control *S. palustris* exhibits during aquatic activity, and further suggests that free-ranging water shrews do not utilize hypothermia to extend underwater foraging endurance. Indeed, our observations of shrews becoming visibly slower and lethargic at T_b’s_ <35°C during the 10-min forced immersion trials in 3°C water implies this level hypothermia would negatively affect foraging performance, though it did not seem to impair their impressive heat generation potential (see below).

Water shrews required to exclusively feed underwater during the 12- to 28-hr trials in 3°C water were observed to consistently elevate their core T_b_ by ∼1.0°C above resting values prior to initiating aquatic foraging bouts. In most cases, this anticipatory increase was associated with increased activity in the terrestrial setup, though it also occurred while the shrews were quietly perched near the water’s edge and likely achieved by a combination of shivering and non-shivering thermogenesis. The latter mode of heat production may be especially critical during and following aquatic foraging bouts, and we have in fact observed large interscapular brown fat deposits in this species (Gusztak, MacArthur, and Campbell, unpublished observations). Notably, free-ranging muskrat and Eurasian otter (*Lutra lutra*) similarly increase T_b_ immediately before initiating foraging bouts in cold water (MacArthur 1979; Kruuk et al. 1987), suggesting this phenomenon may be relatively wide-spread among semi-aquatic mammals.

Possibly owing to the low thermal inertia of water shrews, perhaps in combination with the consumption of cold prey items and the release of peripheral vasoconstriction, T_b_ often declined by up to 1°C following each aquatic excursion in the 24-hr trials (see e.g., Fig. 5). This period was accompanied by vigorous grooming and burrowing in the moss of the chamber such that T_b_ generally returned to 38.5–39.0°C over the ensuing 2-to 4-min recovery period, as was similarly observed in the 20-min behavioral dive trials. Thus, despite their short underwater endurance, minor increases in dive time arising from hypothermia are thus presumably outweighed by the already impressive reaction times afforded by a high T_b_ (20 msec; Catania et al. 2008), which likely enhances foraging efficiency and muscle power output of this diminutive predator. As suggested for free-ranging mink (Harrington et al. 2012), this advantage would be amplified in the physiologically challenging winter season when Q_10_ driven reductions in prey metabolism presumably allows these agile endothermic predators to capture prey items within the short submergence windows (∼10 sec) afforded by their inherently low cADLs. Interestingly, the relatively short (∼65 min) interbout intervals closely align with the mean digesta throughput time of this species (50–55 min; Gusztak et al. 2005), thereby facilitating a dozen or more foraging bouts per day.

### Summary

American water shrews are particularly intriguing from a physiological perspective as allometry predicts this species to have the smallest total body oxygen storage capacity, highest diving metabolic rate, lowest skeletal muscle buffering capacity, and lowest anaerobic capability of any endothermic diver. The water shrew must additionally balance the continual threat of immersion hypothermia while submerged with the pressure to maximize its aerobic dive endurance and anaerobic capacity in order to ensure adequate underwater foraging times. This challenge is presumably greatest in the winter when water shrews must detect and capture prey in near-freezing water, often in total darkness. Surprisingly, the world’s smallest mammalian diver is able to achieve this feat at least in part by elevating core T_b_ by ∼1°C prior to diving. Despite increasing the rate of O_2_ use while underwater, this pre-dive hyperthermia likely enhances their impressive reaction times and set of sensory modalities to efficiently capture energy-rich aquatic prey within time windows as short as 5 to 10 sec. It may also enhance winter foraging efficiency not only by enabling faster underwater swim speeds, but by permitting longer foraging bouts. This pre-dive response may thus provide a critical margin of safety that reduces the risk of costly and potentially debilitating hypothermia.

## Acknowledgments

We thank Chris Schneider, Dean Jeske, Allyson Hindle, and Rob Senkiw for assistance with shrew trapping and husbandry, and Shane Farrow and Stefan Gusztak for lab assistance. Statistical advice and support was kindly provided by Caleb Northam, Jim Hare, and Colin Garroway. R.W.G. was supported in part by a University of Manitoba Faculty of Science Undergraduate Student Research Award, a NSERC Undergraduate Research Award, and a NSERC Postgraduate Scholarship. This research was funded by Natural Sciences and Engineering Research Council (NSERC) Discovery Grants to R.A.M. and K.L.C.

---

**Supplemental Table 1.**
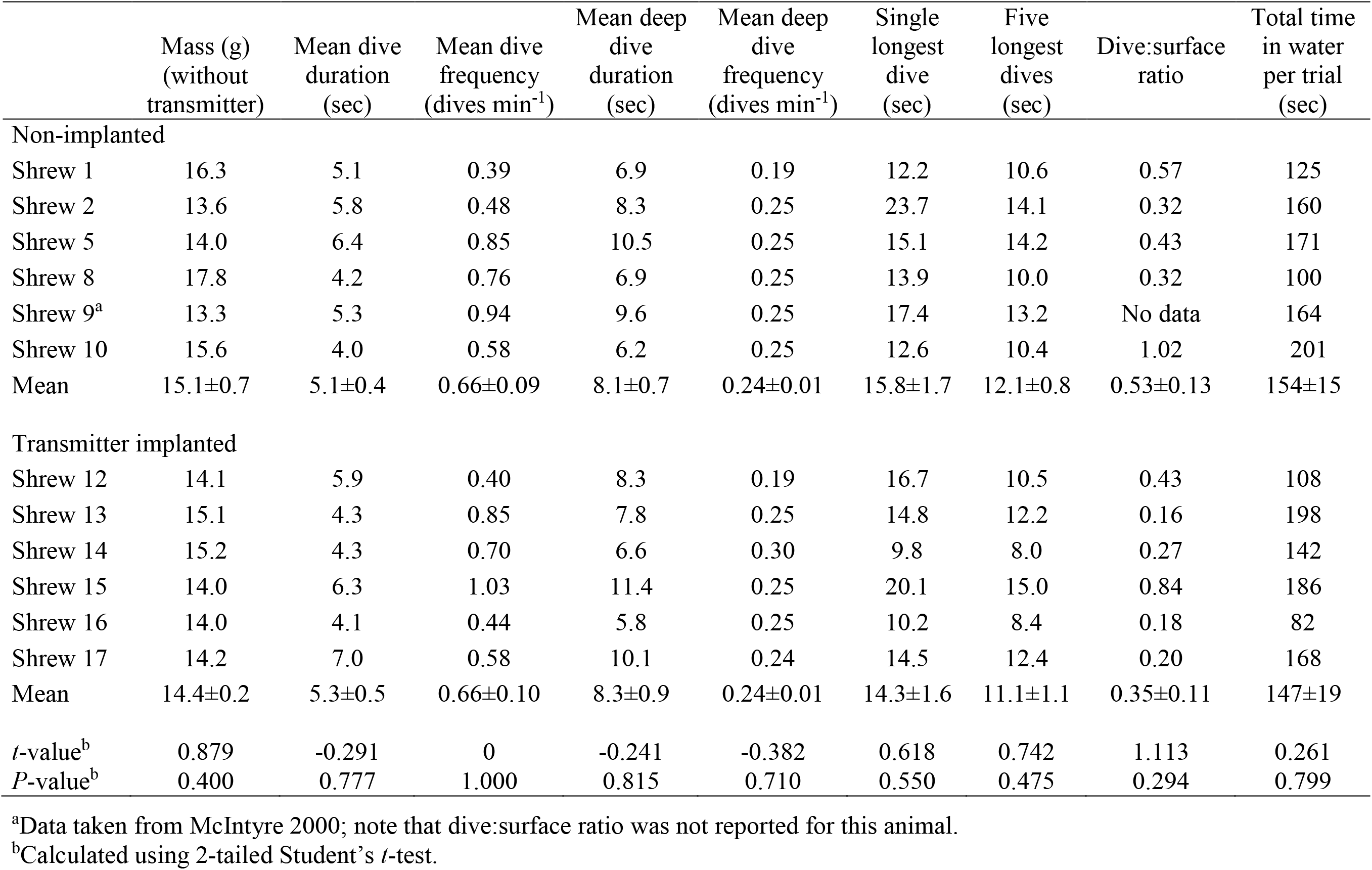
Comparison of the dive performance of adult American water shrews with and without a surgically implanted abdominal temperature transmitter. Dive data for each animal are pooled values from 20-min dive trials completed in 3, 10, 20, and 30°C water.

**Supplemental Table 2.**
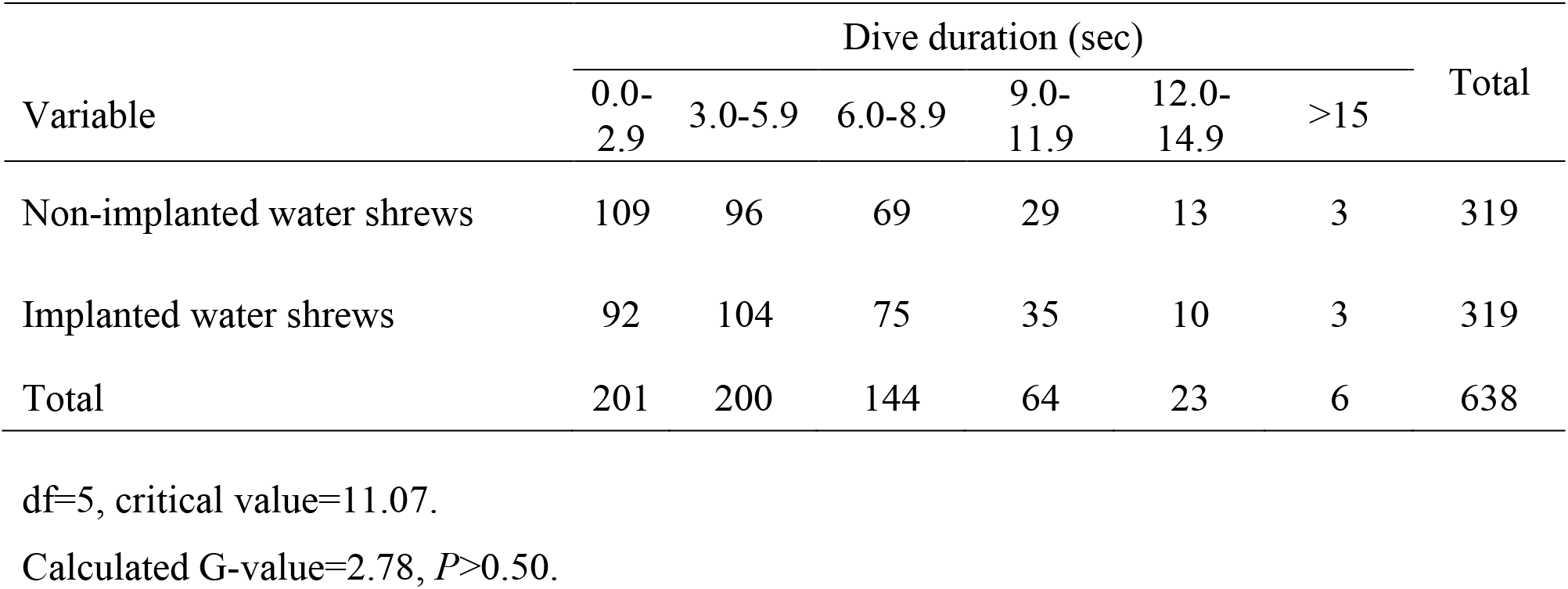
Contingency table using the method of log-likelihood ratio to compare the frequency distribution of voluntary dives by twelve adult American water shrews with (*N*=6) and without (*N*=6) an implanted abdominal temperature transmitter.

**Supplemental Table 3.**
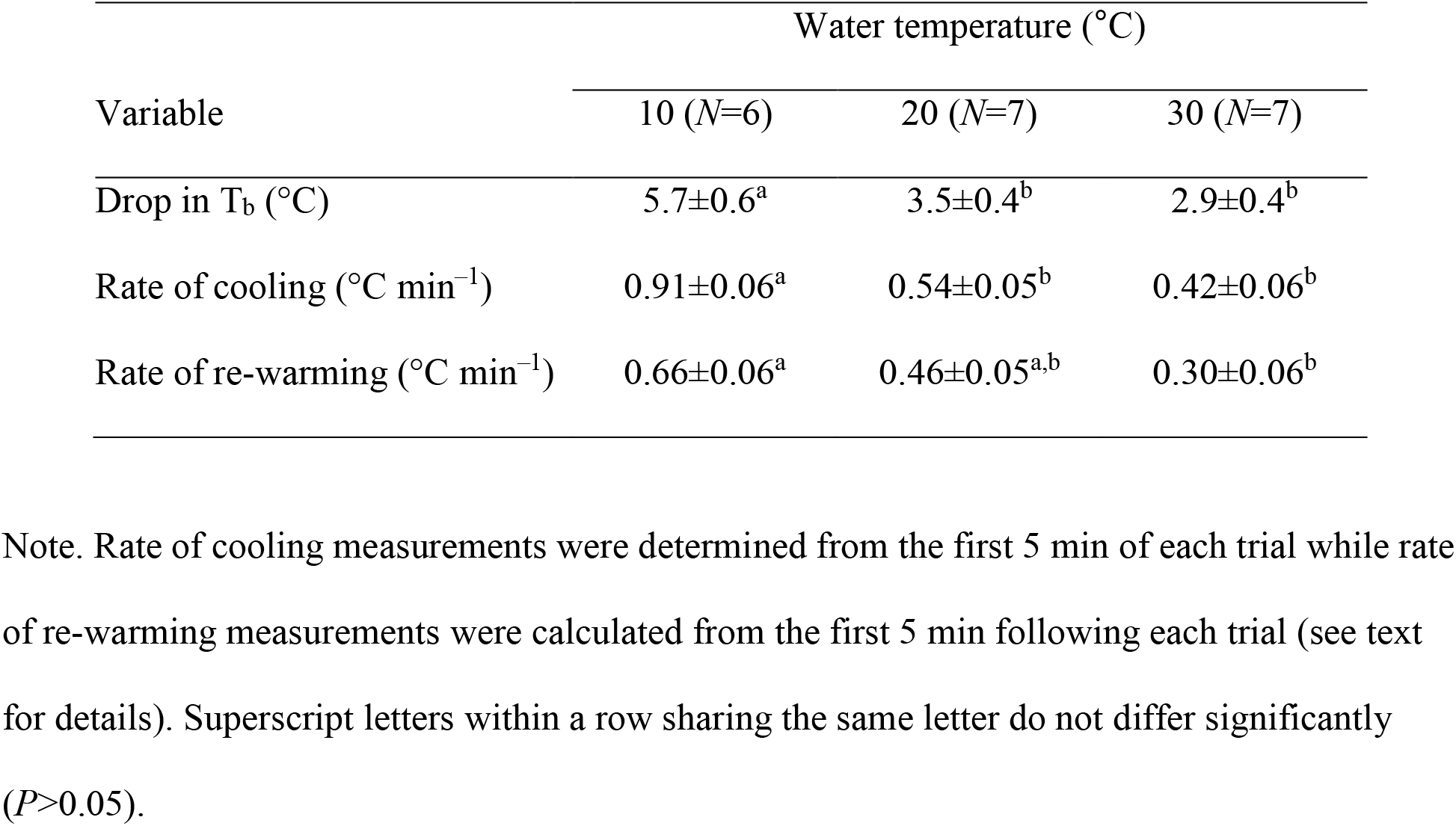
Mean (± 1 SE) body temperature (T_b_) drop, rate of cooling, and rate of rewarming recorded from eight American water shrews during and following 10-min immersion experiments in 10–30°C water. Sample sizes at each water temperature are provided in parentheses.

| Variable | Water temperature (°C) |  |  |
| --- | --- | --- | --- |
| | 10 ($N=6$) | 20 ($N=7$) | 30 ($N=7$) |
| Drop in $T_b$ (°C) | $5.7 \pm 0.6^a$ | $3.5 \pm 0.4^b$ | $2.9 \pm 0.4^b$ |
| Rate of cooling (°C min<sup>-1</sup>) | $0.91 \pm 0.06^a$ | $0.54 \pm 0.05^b$ | $0.42 \pm 0.06^b$ |
| Rate of re-warming (°C min<sup>-1</sup>) | $0.66 \pm 0.06^a$ | $0.46 \pm 0.05^{a,b}$ | $0.30 \pm 0.06^b$ |

Note. Rate of cooling measurements were determined from the first 5 min of each trial while rate of re-warming measurements were calculated from the first 5 min following each trial (see text for details). Superscript letters within a row sharing the same letter do not differ significantly ( $P > 0.05$ ).

**Supplemental Figure 1.**
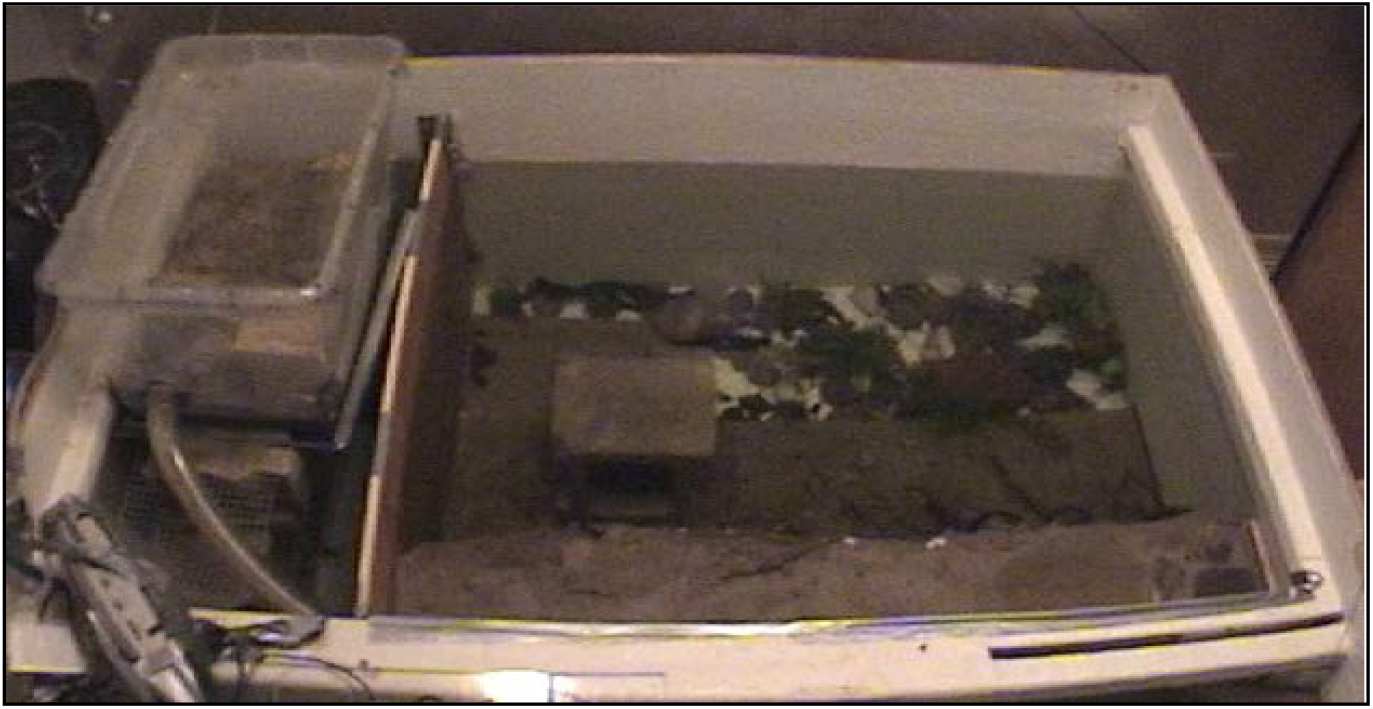
Photograph of the semi-natural riparian set up for observing the voluntary aquatic behavior, body temperature, and activity of five water shrews over periods ranging from 12 to 28 hr.

**Supplemental Figure 2.**
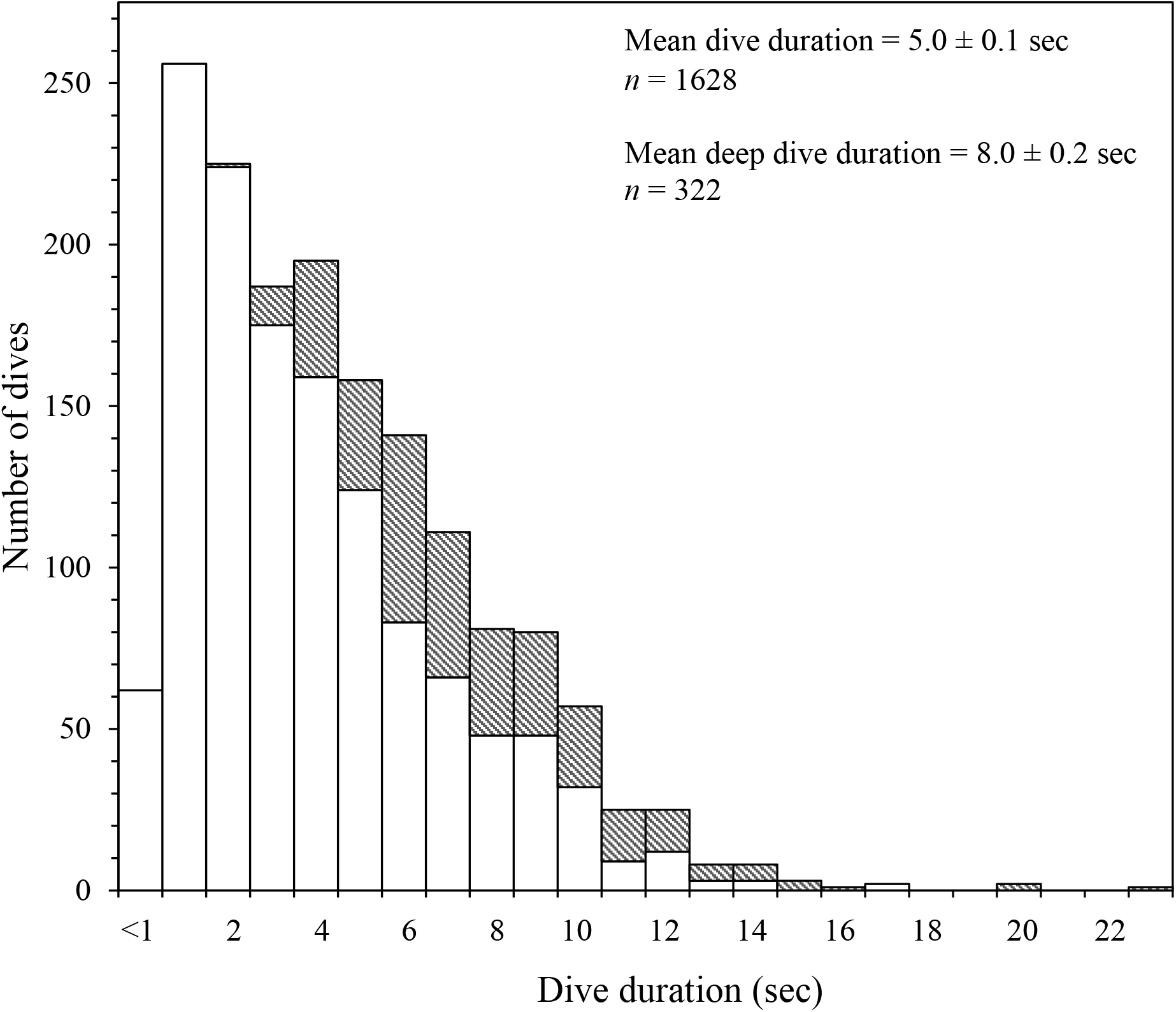
a) Mean number of voluntary dives completed (*n*=317) and b) corresponding mean dive times of 6 transmitter-implanted American water shrews during successive 5-min periods of each 20-min dive trial in 3–30°C water. Bars sharing the same letters are not significantly different (*P*<0.05).

**Supplemental Figure 3.**
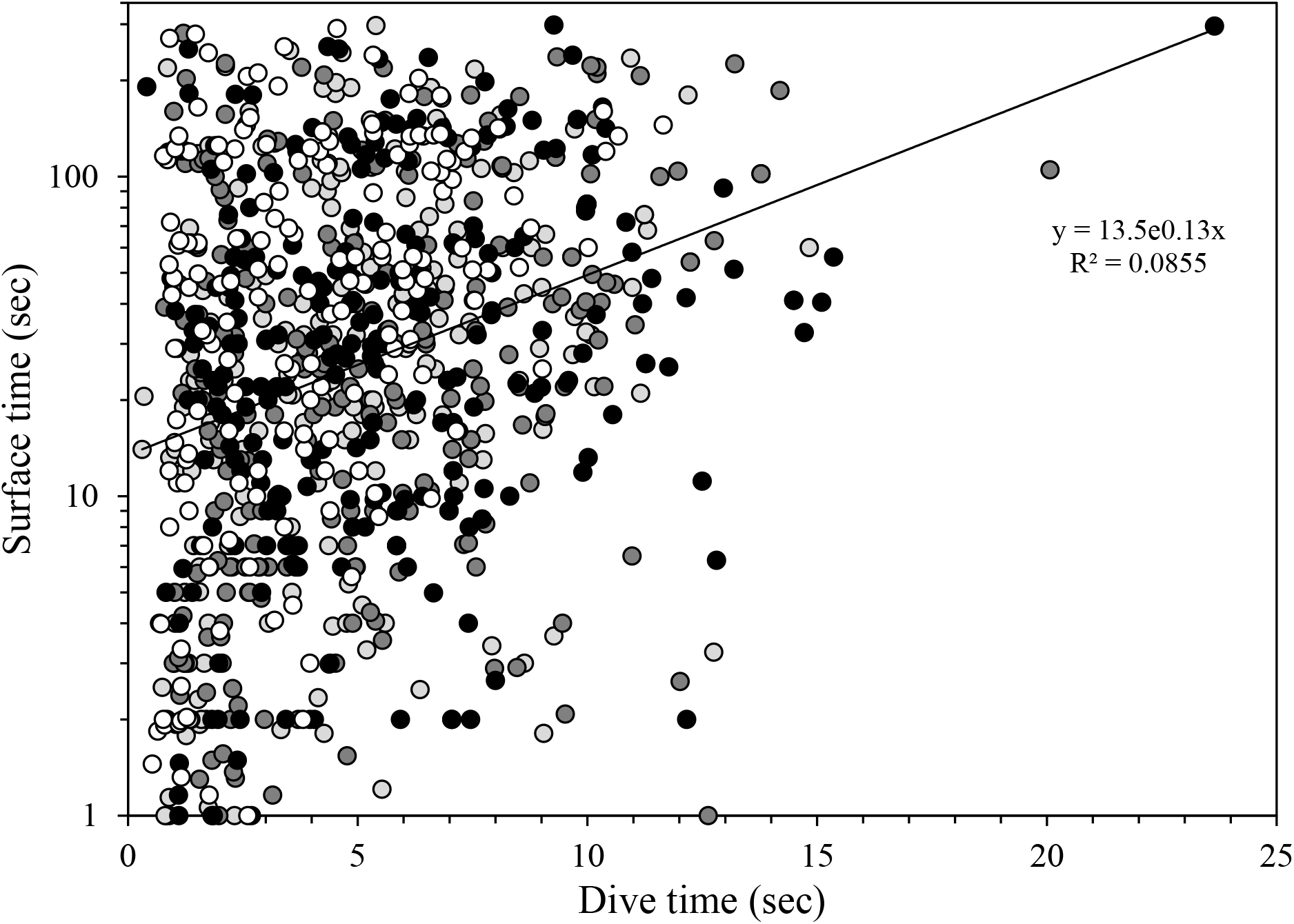
Frequency distribution of voluntary dive times of 25 American water shrews diving in 3–30°C water (*n*=1628 dives). Shallow dives (<10 cm deep) are denoted by open bars, while deep dives (60 cm) are denoted by stippled bars. The mean dive time of all dives combined was 5.0±0.1 sec, while the mean duration of deep dives was 8.0±0.2 sec (*n*=322).

**Supplemental Figure 4.**
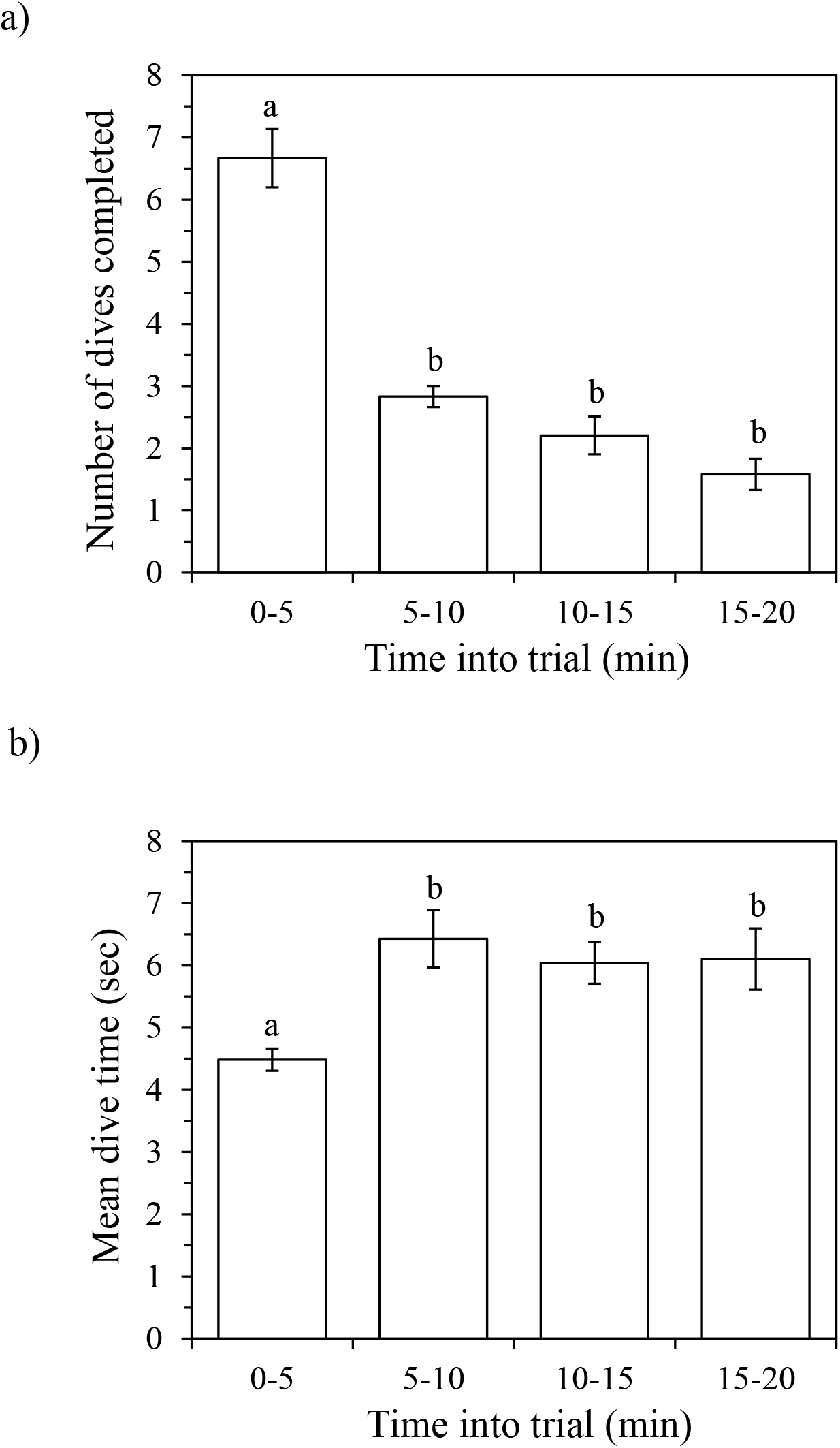
The relationship of inter-dive surface time to duration of preceding dive in water shrews completing consecutive dives during voluntary dive trials. Note, the surface time data was log_10_ transformed to correct for unequal variances. The regression line was fitted by the method of least squares.

**Supplemental Figure 5.**
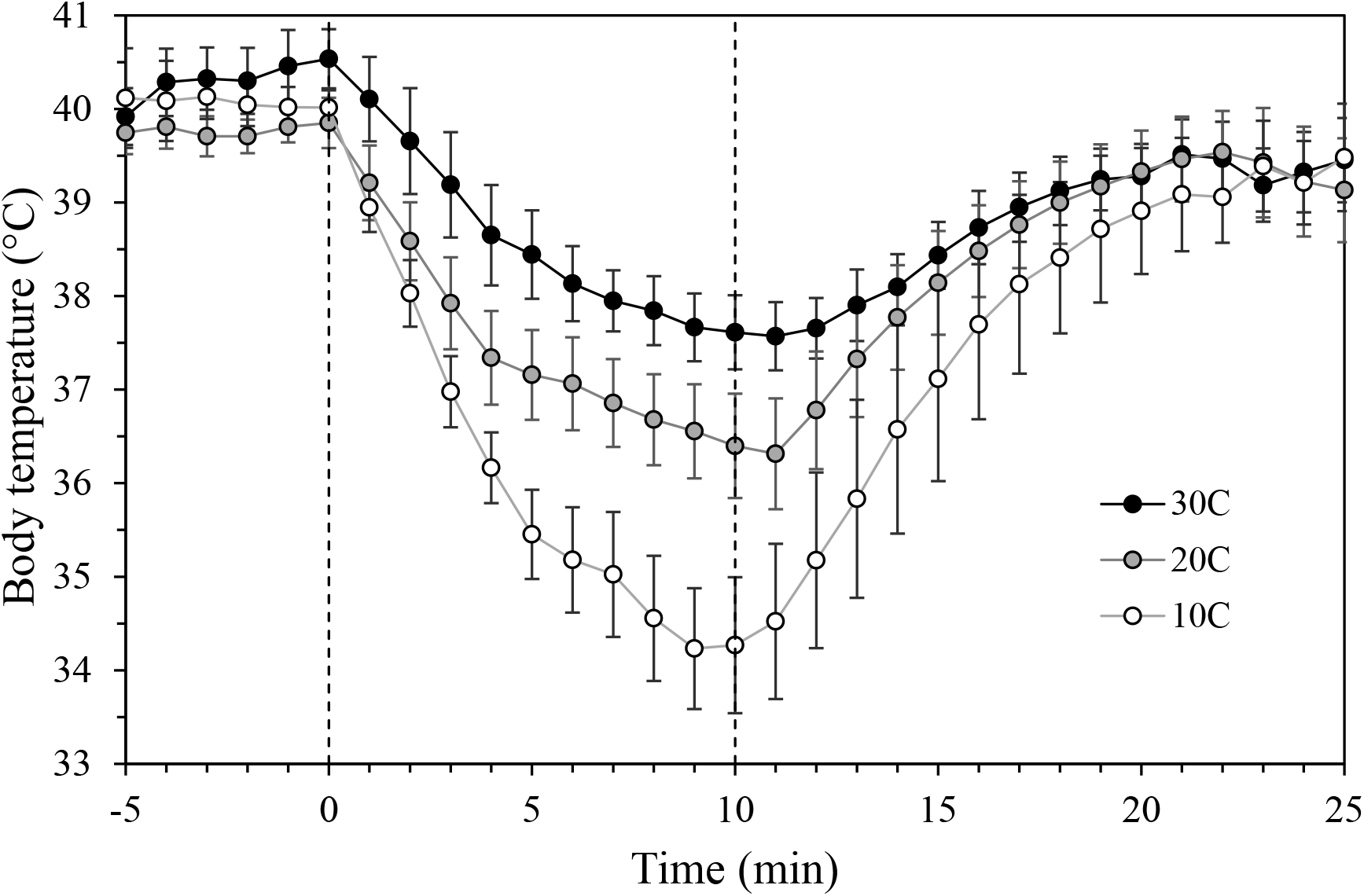
Telemetered body temperature of eight radio-implanted water shrews before and after 10-min immersion trials in 10, 20, and 30°C water.

**Supplemental Figure 6.**
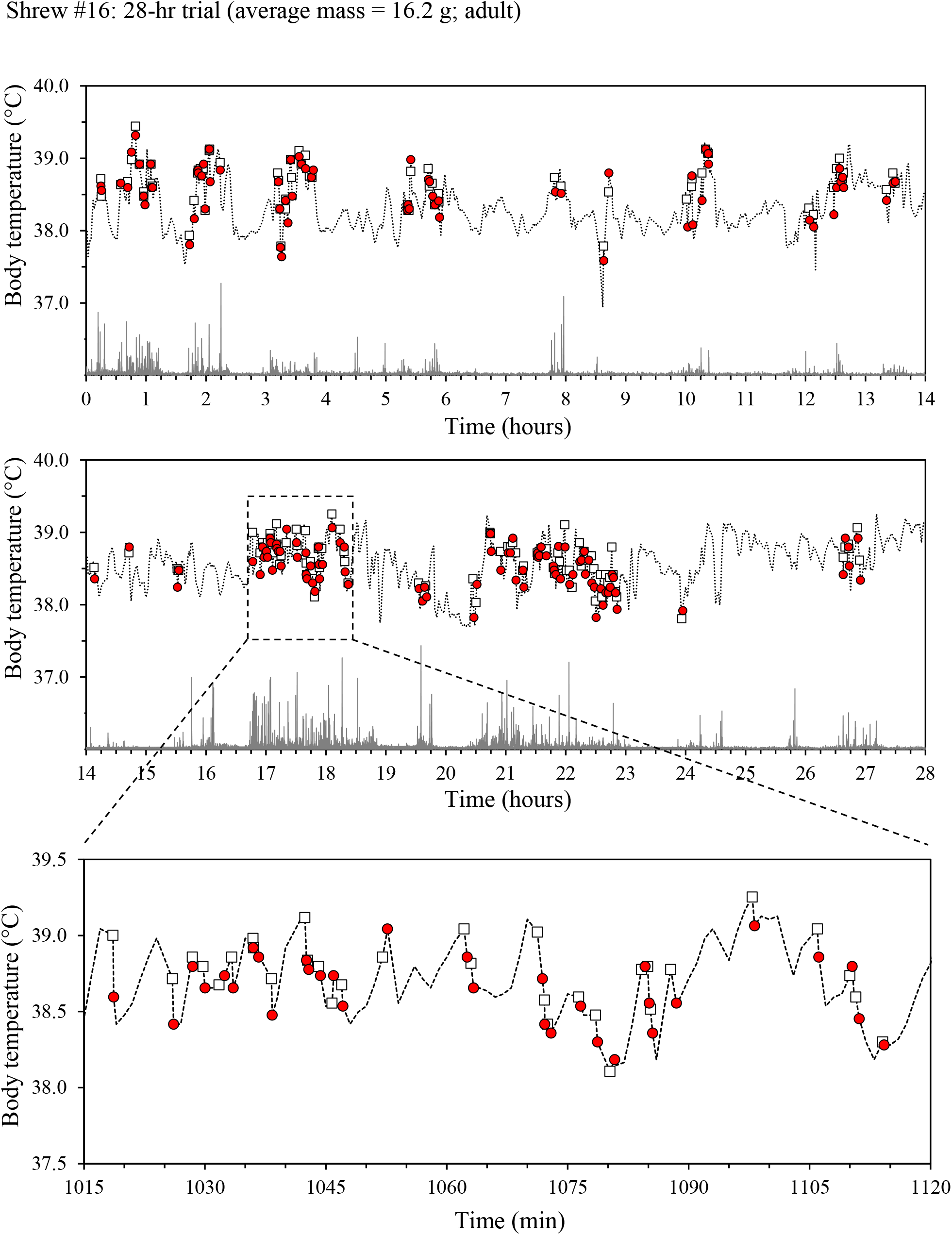
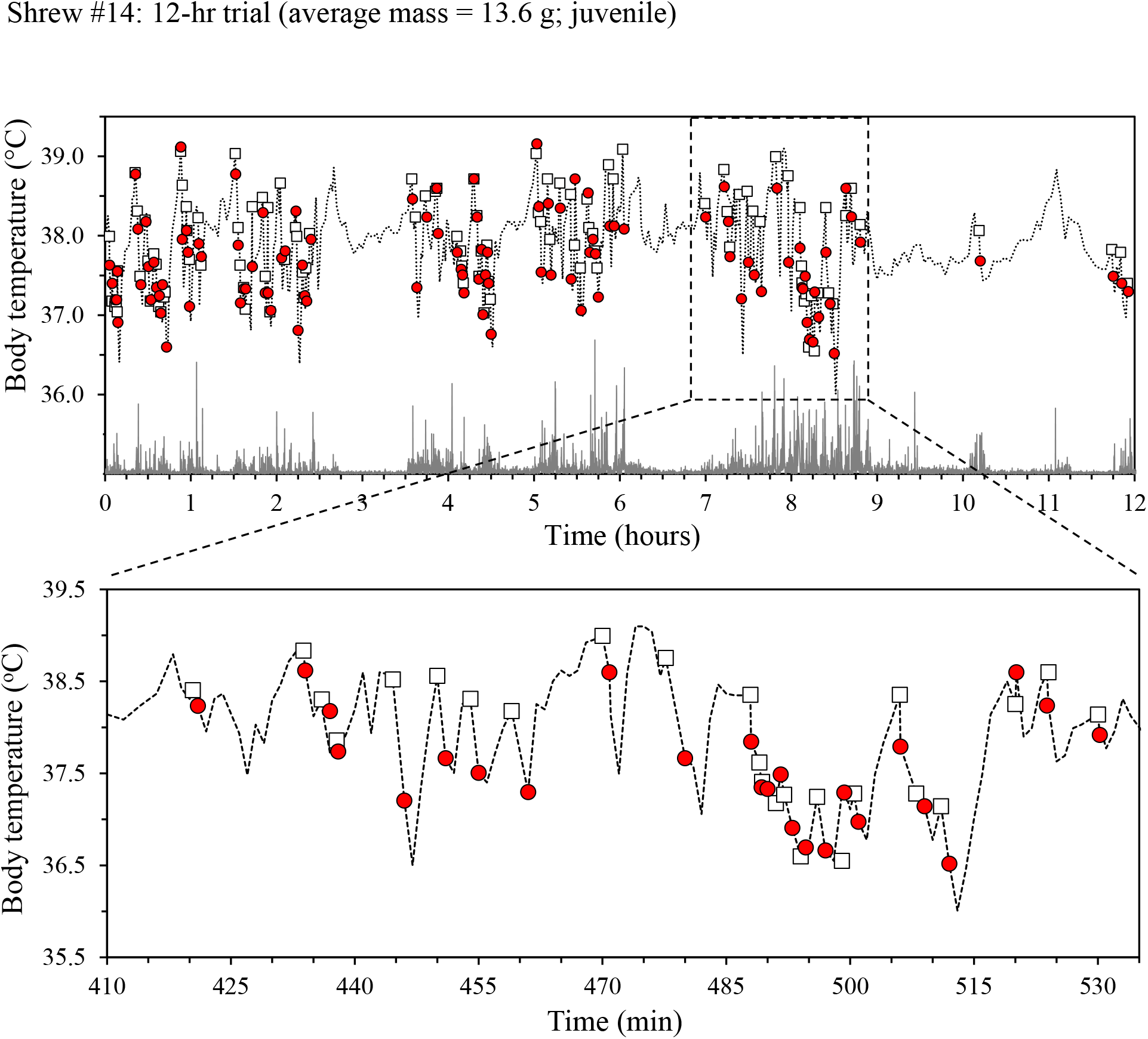
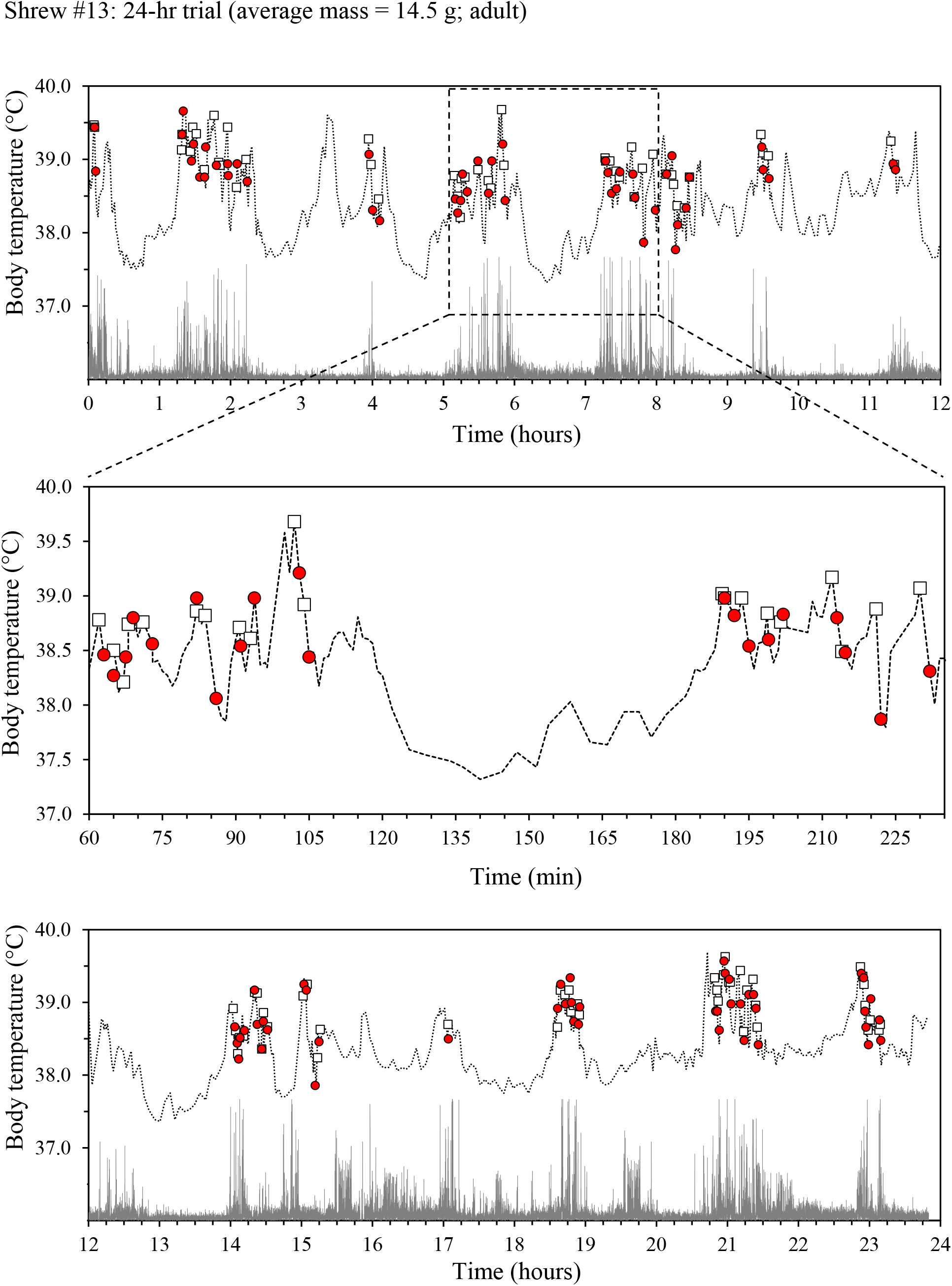
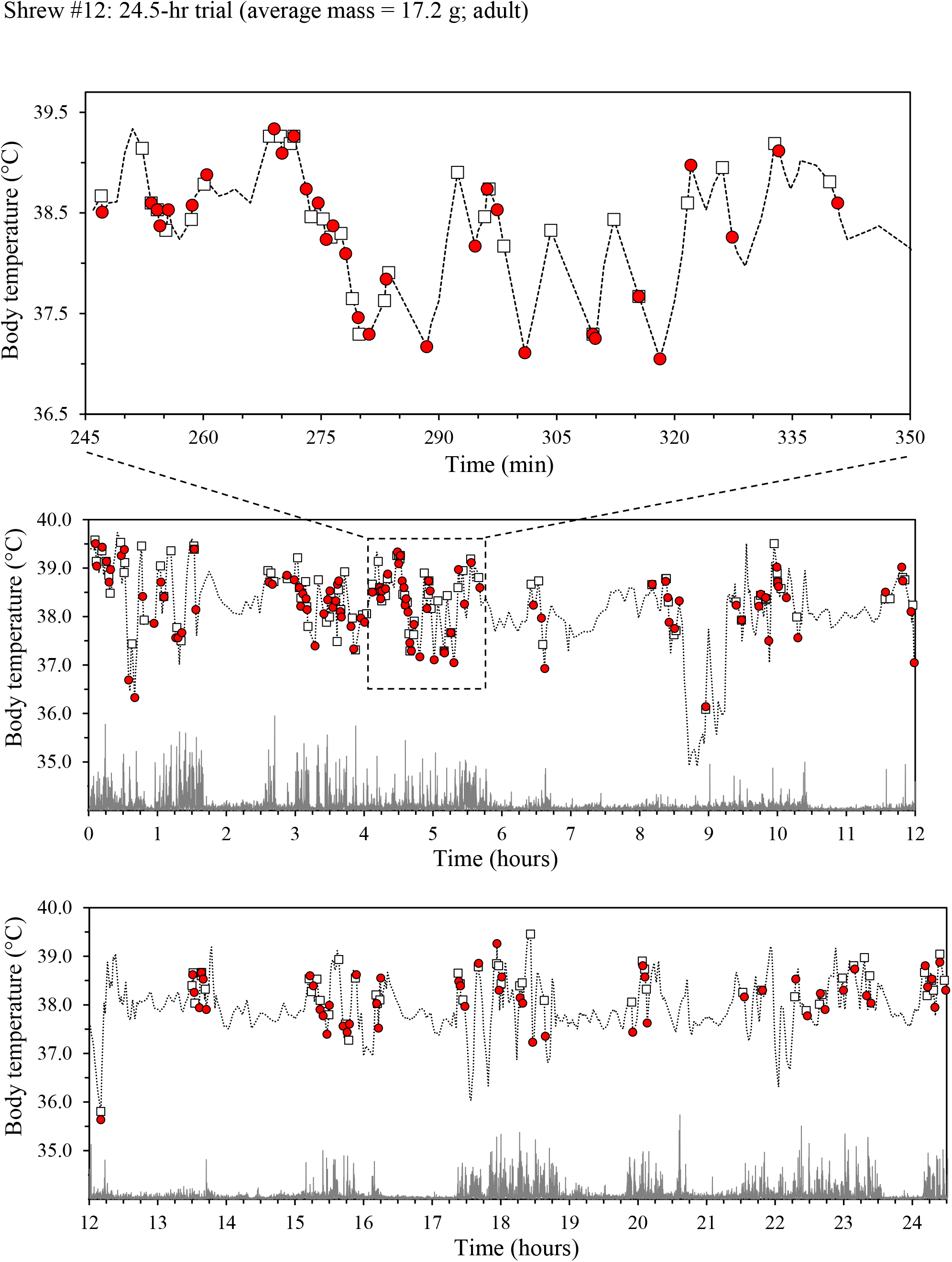
Body temperature fluctuations (dashed lines) and relative activity traces (grey bars) of four American water shrews during 12- to 28-hr trials in a natural riparian environment (water temperature=3°C). Open boxes represent body temperature at the start of each water excursion, while red circles denote body temperature immediately following exit from the water. Expanded inset boxes illustrate fine-scale body temperature profiles over the course of individual dive bouts.

